# Targeting replication stress in neuroblastoma by exploiting the synergistic potential of second generation RRM2 and CHK1 inhibitors

**DOI:** 10.1101/2025.02.26.640375

**Authors:** Iris H. Nelen, Soetkin Leys, Sarah-Lee Bekaert, Fanny De Vloed, Fien Martens, Angeline Praveena Enton Raj, Shunya Ohmura, Lasse Vleminckx, Thomas G. P. Grünewald, Nadine Van Roy, Bram De Wilde, Frank Speleman, Annelies Van Hemelryk, Lisa Depestel, Kaat Durinck

## Abstract

Tumor cells often cope with elevated levels of replication stress (RS) causing increased dependency on ATR-CHK1 signalling. We previously presented RRM2, the regulatory component of the ribonucleotide reductase (RNR) enzyme, as novel dependency in neuroblastoma, in keeping with its role in RS resistance. We identified strong synergism for combined RRM2-CHK1 inhibition using the iron chelator triapine and prexasertib respectively. To obtain direct RNR targeting, we evaluated a novel inhibitor, TAS1553, specifically disrupting the RNR complex in this study. Treatment with TAS1553 impedes cell growth and induces enhanced RS, DNA damage and apoptosis. We demonstrated strong synergism between TAS1553 the CHK1 inhibitors prexasertib and SRA737 in both NB and sarcoma cell lines, underscoring the broad clinical potential of combinatorial RRM2-CHK1 inhibition. Transcriptome profiling demonstrated strong overlap between the different RRM2-CHK1 treatments and revealed differential expression of RNA splicing components, opening new perspectives for combination treatments using splicing inhibitors. Altogether, this study paves the way for further preclinical testing of second generation RRM2 and CHK1 inhibitors such as TAS1553 and SRA737 in neuroblastoma and sarcomas.

## Introduction

Cancer cells often display high proliferation rates causing increased levels of replication stress (RS), defined as slowing, stalling or the collapse of replication forks, resulting in increased DNA damage (Zeman & Cimprich, 2014). Neuroblastoma (NB), a deadly pediatric tumor of the sympathetic nervous system, displays high basal levels of RS (Cole *et al*, 2011) and adopts a RS resistance phenotype, which renders them particularly sensitive for pharmacological inhibition of the ATR-CHK1 replicative stress induced signaling pathway (Durinck & Irwin, 2024). Indeed, previous studies indicate that NB is one of the most sensitive tumors to CHK1 inhibition (Cole *et al*, 2011; Blosser *et al*, 2020). Moreover, studies in ALK-driven genetically engineered mouse models of NB showed complete tumor regression upon ATR inhibition, highlighting the effectiveness of therapeutically targeting the DNA damage response (DDR) pathway in NB (Szydzik *et al*, 2021; Borenäs *et al*, 2024).

Cancer cells exhibiting enhanced RS are particularly vulnerable to dNTP depletion during early S-phase and therefore require continuous and sufficiently high levels of dNTP pool supplementation. In line with this observation, we previously identified ribonucleotide reductase subunit M2 (RRM2) as a novel therapeutic target, which is synergistic with pharmacological CHK1 inhibition in high-risk NB (Nunes *et al*, 2022). RRM2 is part of the ribonucleotide reductase (RNR) enzyme, where it forms a heterotetrametric dimer together with RRM1, which catalyses the rate-limiting step in dNTP synthesis (Aye *et al*, 2014). In contrast to RRM1, RRM2 expression is tightly regulated during cell cycle progression with elevated levels during the S-phase (Chabes & Thelander, 2000; D’Angiolella *et al*, 2012). The functional importance of RRM2 is more broadly marked by its overexpression in several adult cancers, including lung cancer (Bokhari *et al*, 2022; Jin *et al*, 2020), prostate cancer (Mazzu *et al*, 2019) and melanoma (Granieri *et al*, 2022; Fatkhutdinov *et al*, 2016). More recent work has also converged towards its critical role also in the context of pediatric entities including Ewing sarcoma (EWS) (Koppenhafer *et al*, 2020; Ohmura *et al*, 2021), retinoblastoma (Yang *et al*, 2021), atypical teratoid rhabdoid tumors (Giang *et al*, 2024, 2023) and hepatoblastoma (Brown *et al*, 2023).

Given the broad role of RRM2 across multiple tumor types, several RRM2 inhibitors have been developed, such as hydroxyurea (HU) and triapine (3-AP). HU penetrates the RNR complex and destroys the tyrosyl free radical via one electron oxidation (Atkin *et al*, 1973; Lassmann *et al*, 1992). Consequently, the nucleoside diphosphate (NDP) reduction reaction is inhibited. 3-AP is an iron chelator, interacting with the iron molecule used by RRM2. This interaction results in the production of reactive oxygen species that inactivate the RNR complex via quenching of the tyrosyl free radical (Shao *et al*, 2006). Both compounds have been tested extensively in cancer patients but were shown to be associated with rapid resistance and significant toxicity (Knox *et al*, 2007; Zhan *et al*, 2021).

Protein-protein interaction (PPI) inhibitors are an emerging topic in the field of drug development, including for targeting the DDR pathway, as illustrated by compounds disrupting the BRCA2-RAD51 and the MUS81-EME1 interactions (Scott *et al*, 2021; Zhang *et al*, 2023). Recently, Ueno et al. developed TAS1553, a PPI disruptive agent targeting the RNR complex. TAS1553 binds to the C-terminus binding site of RRM1, thereby inhibiting the physical binding of RRM2 (Ueno *et al*, 2022). Given this specific mode of action, TAS1553 provides a unique entry point for more selective and direct RNR inhibition compared to the historical set of available drugs and could potentially reduce the resistance and toxicity observed using HU and 3-AP.

In the current study, we show that TAS1553 inhibits NB cell proliferation and induces RS, DNA damage and apoptosis. We benchmarked the molecular responses of TAS1553 single drug treatment and revealed a strong impact on RNA splicing. Second, in line with our previous work (Nunes *et al*, 2022), we also demonstrate strong synergism between TAS1553 and the CHK1 inhibitors prexasertib and SRA737 in NB cell lines, with underlying gene signatures recapitulating our previously established gene signatures (Nunes *et al*, 2022). We confirmed these synergistic interactions *in vivo* using a zebrafish neuroblastoma xenograft model and expanded the validity of these drug combinations *in vitro* towards pediatric sarcoma.

## Results

### TAS1553 mediated RRM1-RRM2 disruption inhibits neuroblastoma cell growth and induces replicative stress, DNA damage and apoptosis

The phenotypic impact of TAS1553 treatment was assessed using a broad concentration range (0.5µM – 4µM) in a panel of NB cell lines including both *MYCN* amplified and non-amplified tumors. TAS1553 treatment induced a decrease in cell confluence in eight out of ten cell lines with IC_50_-values in the low micromolar range (**Fig 1A**), with no significant differential response profile between *MYCN* amplified and non-amplified cell lines. In addition, TAS1553 also inhibits cell growth in a panel of pediatric sarcoma cell lines, including EWS, osteosarcoma and rhabdomyosarcoma (**Appendix Fig S1**). Furthermore, we observed a cytotoxic effect of TAS1553 in NB cell lines through apoptotic cell death compared to DMSO controls, while only cytostatic effects were noted in the non-cancerous retinal pigment epithelial (RPE) cells, underscoring the on-target effect of TAS1553 in cancer cell lines (**Fig 1A, B**). In line with our previous findings regarding RRM2 inhibition using 3-AP in NB (Nunes *et al*, 2022), TAS1553 treatment resulted in increased RS levels and both single and double stranded DNA breaks, represented by increased levels of ATR induced pCHK1^S345^, pRPA32^S33^, and *γ*H2AX (**Fig 1C**). We also noted a slight upregulation of RRM2 levels upon TAS1553 treatment (**Fig 1C**), which could point towards a compensatory upregulation of RRM2, as previously also reported for Glioblastoma upon RRM2 inhibition (Corrales-Guerrero *et al*, 2023). Furthermore, induction of apoptosis was confirmed through the elevated protein levels of cleaved PARP (cPARP) and increased mRNA expression of the pro-apoptotic genes *BAX*, *PUMA* and *NOXA* (**Fig 1C, D)**. The expression of the p53 target genes *CDKN1A* and *RRM2B* was also upregulated following TAS1553 treatment (**Fig 1D**). Functional inhibition of the RNR complex was validated by depletion of the dNTP pools after TAS1553 treatment (**Fig 1E**). Altogether, our *in vitro* phenotypic response data support that TAS1553 is effective in suppressing NB and sarcoma cell growth by enhancing RS, increasing DNA damage and inducing cell death.

**Figure 1:**
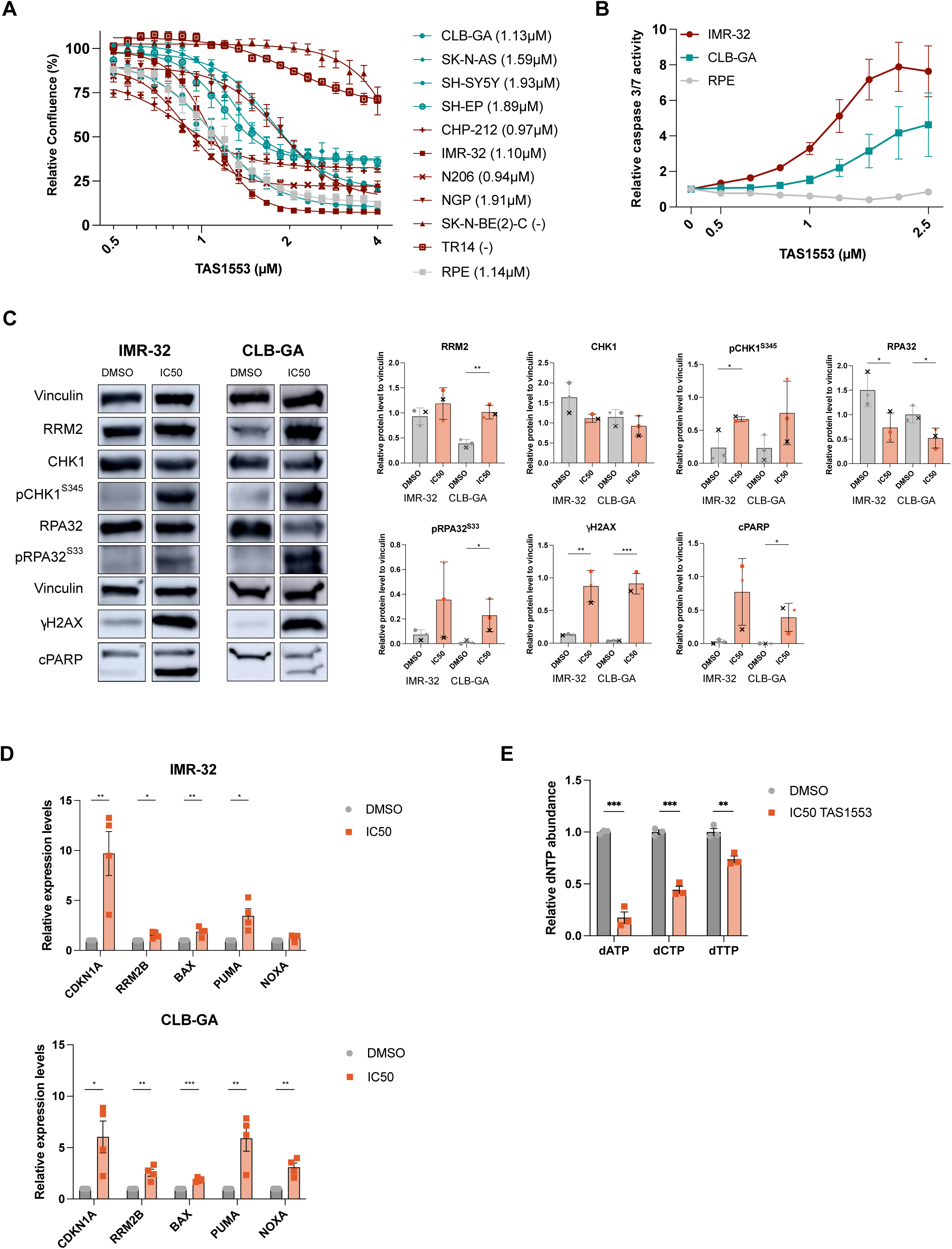
TAS1553 reduces neuroblastoma cell growth and induces RS stress, DNA damage and apoptosis *in vitro* and *in vivo*. A. Dose-response curves following 96h of TAS1553 exposure in a panel of neuroblastoma cell lines, including 6 *MYCN* amplified (*red*) and 4 *MYCN* non-amplified (*blue*) lines, and the nonmalignant RPE cell line (*grey*) using the IncuCyte® Live-Cell imaging. The IC_50_ of each cell line is indicated between the brackets. Graphs represent the mean of 3-5 biological replicates in triplicate ± standard error of the mean (SEM). B. Caspase-Glo luminescence-based measurements of caspase 3/7 activity in IMR-32, CLB-GA and RPE following TAS1553 treatment for 48h. Graphs represent the mean of 3 biological replicates in triplicate ± SEM. C. Representative immunoblotting data for various replicative stress and DNA damage markers in IMR-32 and CLB-GA following 48h treatment with their respective IC_50_ of TAS1553 (1.18µM and 1.54µM respectively) (*left*). Quantification of the immunoblotting data obtained for three independent biological replicates (indicated by symbols) (*right*). Significance is measured via an unpaired T-test (*= p≤0.05, **= p≤0.01), error bars indicate standard deviation (SD). D. RT-qPCR based measurements of mRNA expression levels of the apoptotic genes *BAX, PUMA* and *NOXA*, and the *p53* target genes *CDKN1A* and *RRM2B* after treatment of IMR-32 (*top*) and CLB-GA (*bottom*) cell lines with their respective IC_50_ of TAS1553 at 48h. Graphs represent the mean of 4 biological replicates ± SEM. *p≤0.05, **p≤0.01, ***p≤0.001 (multiple unpaired t-test). E. dNTP pools are reduced in IMR-32 upon treatment with the IC_50_ of TAS1553 for 48h. Significance was measured using the unpaired T-test (**= p≤0.01, ***= p≤0.001), error bars represent SEM.

### TAS1553 treatment induces replication stress, apoptosis and immunomodulatory signatures in neuroblastoma

Transcriptome profiling following TAS1553 exposure revealed a robust set of differentially expressed genes in both the MYCN amplified as well as non-amplified cancer cell lines IMR-32 and CLB-GA respectively (**Fig 2A and Appendix Fig S2A**). Interestingly, one of the top ranked downregulated genes is *H1-0*, encoding the linker histone H1, with a key role in replication fork stability upon fork stalling (Ozgencil *et al*, 2023). Reduced expression of *H1-0* under therapeutic pressure suggests increased fork stalling and possible fork collapse, which induces cell death. Furthermore, genes involved in apoptosis (*CDKN1A, BBC3, TNFRSF10B*) and DNA damage (*PLK2, GADD45A*) were amongst the top-ranked upregulated genes in both IMR-32 and CLB-GA (**Fig 2A**). Gene set enrichment analysis (GSEA) revealed on the one hand a significant upregulation of p53 target genes, including *HEXIM1* as a negative regulator of the transcriptional regulator pTEFb (Tan *et al*, 2016) (**Fig 2B, C**). Moreover, upregulation of TNF signaling via NFκB and interferon alpha associated gene signatures can potentially be linked to an immunomodulatory drug response. On the other hand, MYC target genes, such as *CBX3*, implicated in replication fork stability (Ma et al, 2024), and G2M checkpoint related genes, including the NB core regulatory circuitry factor *MEIS2* (De Wyn *et al*, 2021), were amongst the top downregulated gene sets in both IMR-32 and CLB-GA (**Fig 2B, C, Appendix Fig S2B**), in accordance with our previously established 3-AP treated or siRNA mediated RRM2 knock-down transcriptome signatures in NB (Nunes *et al*, 2022) (**Fig 2B, Appendix Fig S2C**). Given that the clinical trials for 3-AP as RRM2 inhibitor were associated with major adverse effects, including methemoglobinemia, we propose TAS1553 as a potent alternative to be further investigated in extended preclinical testing and subsequent clinical trials.

**Figure 2:**
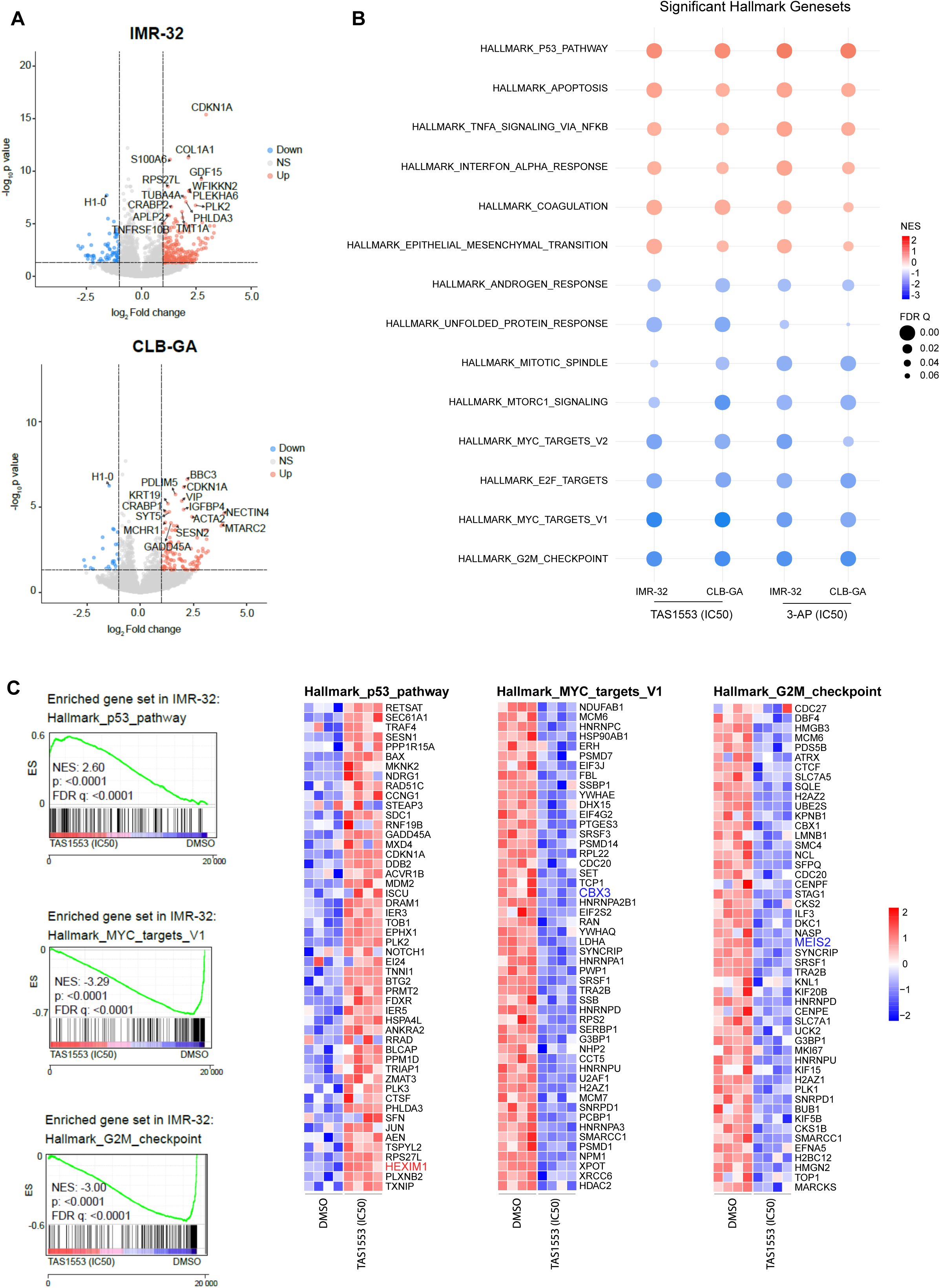
Benchmarking TAS1553 imposed transcriptional signatures in neuroblastoma. A. Volcano plots of IMR-32 (*top*) and CLB-GA (*bottom*) of significantly differentially expressed genes (DEGs) following TAS1553 IC_50_ exposure for 48h (1.18µM and 1.54µM respectively). Genes indicated by name cover the top 15 DEGs. B. Bubble plots of all overlapping Hallmark MSigDB gene sets significantly up or downregulated in IMR-32 or CLB-GA cells after treatment (48h) with their respective IC_50_ for TAS1553 (1.18µM and 1.54µM) (*left*) or 3-AP (0.69µM and 0.96µM) (*right*). C. Detailed view of the GSEA enrichment for the p53 pathway, MYC targets and G2/M checkpoint Hallmark MSigDB gene sets after treatment of IMR-32 with 1.18µM TAS1553 (IC_50_) treatment (*left*) and heatmaps showing the top 50 genes within the core enrichment (*right*). ES = enrichment score.

### Benchmarking the TAS1553-prexasertib drug synergism *in vitro* and *in vivo*

Following the validation of TAS1553 as a potent RRM2 inhibitor, we next evaluated whether TAS1553 treatment in combination with CHK1 inhibition using prexasertib also resulted in similar synergy as previously obtained with 3-AP (Nunes *et al*, 2022). To this end, we tested a broad concentration range of both compounds (0.06µM-1.50µM of TAS1553 and 0.1nM-2nM of prexasertib) and monitored cell confluence for 96 hours via the IncuCyte Live Cell Imaging System (**Fig 3A, Appendix Fig S3A)**. Treatment with 0.88µM TAS1553 and 0.74 nM prexasertib resulted in high bliss index (BI) scores of respectively 0.80 and 0.66 in IMR-32 and CLB-GA, indeed confirming strong synergism between TAS1553 and prexasertib. This combination treatment resulted in reduced cell confluence and increased caspase 3/7 levels reflecting apoptosis, compared to treatment with the same low-dose concentrations of the single agents or DMSO control treatment (**Fig 3A, Appendix Fig S3A**). In the non-cancerous RPE cells, only a slight reduction in cell confluence and no apoptotic effects were observed, suggesting on-target toxicity (**Fig 3B**). In line with the increased caspase 3/7 activity in IMR-32 and CLB-GA, mRNA expression of the pro-apoptotic genes *BAX*, *PUMA* and *NOXA* was upregulated following combination treatment while increased gene expression of *CDKN1A* and *RRM2B* indicates p53 pathway activation (**Fig 3C, Appendix Fig S3C**). Immunoblotting showed elevated pCHK1^S345^, pRPA32^S33^ and *γ*H2AX levels, indicating increased RS levels and DNA damage induction. In addition, we observed downregulation of pCHK1^S296^ (CHK1 autophosphorylation site) and total CHK1 levels, in line with previous studies showing that functional inhibition or knockdown of RRM2 directly affects protein translation through inhibition of the translational inhibitor 4E-BP1 (Goss *et al*, 2021). Furthermore, elevated cPARP levels confirmed the observed apoptosis induction in NB cells treated with the combination compared to single agent or DMSO control treatment conditions (**Fig 3D, Appendix Fig S3D**). Next, we also confirmed this synergistic interaction in a panel of pediatric sarcoma cell lines including one EWS, one osteosarcoma and two rhabdomyosarcoma lines (**Appendix Fig S4A-C**), showing therapeutic value for this combination treatment in other pediatric entities beyond NB.

**Figure 3:**
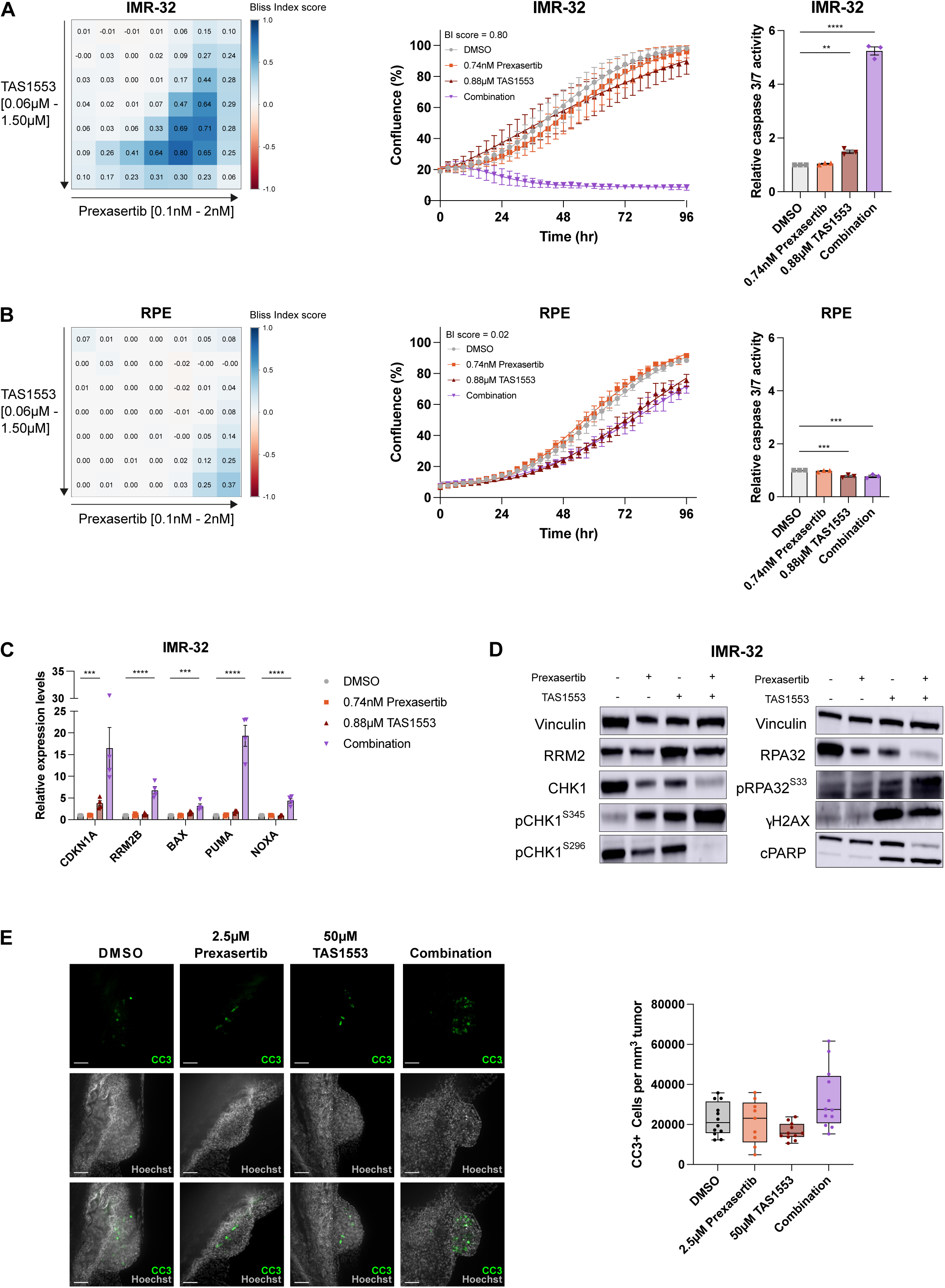
Benchmarking TAS1553-prexasertib drug synergism *in vitro* and *in vivo*. A. Heatmap showing the Bliss Index (BI) scores for IMR-32 after 96h of treatment with a range of concentrations of TAS1553 and prexasertib. Biological replicates= 4 in duplicate (*left*). Cell confluency was monitored for 96h and cells were treated with the concentrations with the highest BI score (0.74nM prexasertib and 0.88µM TAS1553). Biological replicates= 4 in duplicate, error bars represent SEM (*middle*). Caspase-Glo luminescence-based measurements of caspase 3/7 activity in IMR-32 following treatment with TAS1553 and prexasertib for 48h. Biological replicates= 3 in triplicate, error bars represent SEM. Significance was measured using one-way ANOVA followed by a Dunnett’s multiple comparison test (**= p≤0.01, ****= p≤0.0001) (*right*). B. Heatmap showing the BI scores for RPE after 96h of treatment with a range of concentrations of TAS1553 and prexasertib. Biological replicates= 3 in duplicate (*left*). Cell confluency was monitored for 96h and cells were treated with the concentrations with the highest BI score in NB (0.74nM prexasertib and 0.88µM TAS1553). Biological replicates= 3 in duplicate, error bars represent SEM (*middle*). Caspase-Glo luminescence-based measurements of caspase 3/7 activity in RPE following treatment with TAS1553 and prexasertib for 48h. Biological replicates= 3 in triplicate, error bars represent SEM. Significance was measured using one-way ANOVA followed by a Dunnett’s multiple comparison test (***= p≤0.001) (*right*). C. RT-qPCR based measurements of mRNA expression of apoptotic genes *BAX, PUMA* and *NOXA*, and the *p21* target genes *CDKN1A* and *RRM2B* after treatment of IMR-32 with the combination of TAS1553 (0.88µM) and prexasertib (0.74nM) that resulted in the highest synergy scores at 48h. Biological replicates= 4 in duplicate. Significance was measured using the one-way ANOVA followed by a Dunnett’s multiple comparison test (*= p≤0.05, ***= p≤0.001, ****= p≤0.0001), error bars represent SEM. D. Representative immunoblotting data for various replicative stress and DNA damage markers in IMR-32 following 48h treatment with 0.74nM prexasertib and 0.88µM TAS1553 for 48h. The quantification of all replicates (n = 3) is shown in Supplementary Figure S5A. E. Representative image of zebrafish treated with DMSO or 2.5µM prexasertib and 50µM TAS1553. CC3 is shown in green and Hoechst in grey, scale bar represents 50µm (*left*). Boxplot showing increased cleaved caspase 3 (CC3+) expression in SK-N-AS NB cells injected in zebrafish after 96h of treatment with 2.5µM prexasertib and 50µM TAS1553 (*right)*.

To assess the *in vivo* efficacy of combined TAS1553 and prexasertib treatment, we utilized a zebrafish NB xenograft model using the SK-N-AS cells, given its well-established engrafting potential (Lawrence *et al*, 2024). First, the maximum non-toxic dose (MNTD) was determined by testing a concentration range of TAS1553 and prexasertib on wild-type 3-days-old zebrafish. Zebrafish tolerated 50µM of TAS1553 and 2.5µM prexasertib without any apparent signs of toxicity (cardiac oedema or curved tails) (**Appendix Fig S5A**). Next, SK-N-AS cells were injected into the perivitelline space of 2-days-old zebrafish, and treated with the respective MNTD of both compounds 24 hours after engraftment. The compounds were refreshed every 24 hours for a total treatment period of 96 hours. Zebrafish xenografts treated with the combination show increased levels of cleaved caspase 3 (CC3+), a marker for apoptosis, compared to single agent and DMSO control treated zebrafish (**Fig 3E**), in line with our *in vitro* findings.

### Expanding the combinatorial RRM2-CHK1 drugging with TAS1553 and the selective CHK1 inhibitor SRA737

Given that prexasertib also substantially inhibits CHK2 (King *et al*, 2015), we next assessed combinatorial treatment of TAS1553 with SRA737, which has been shown to target CHK1 with a 1,000-fold greater selectivity over CHK2 (Osborne *et al*, 2016). We observed higher BI scores for TAS1553-SRA737 drugging (up to 0.84 in IMR-32 and 0.70 in CLB-GA) (**Fig 4A, Appendix Fig 3B**), compared to the combination of TAS1553 and prexasertib (0.80 and 0.66 in IMR-32 and CLB-GA respectively). Moreover, these high BI scores were achieved with lower concentrations of TAS1553 (0.88µM in combination with prexasertib versus 0.51µM in combination with SRA737). These strong synergistic effects were again confirmed in a panel of pediatric sarcoma cell lines of EWS, osteosarcoma and rhabdomyosarcoma, while not affecting the non-cancerous RPE cells, showing on-target toxicity (**Fig 4B, Appendix Fig S4D-F**). The reduced confluency was again accompanied by an increase in apoptosis, specifically in the cancer cell lines and not in the non-cancerous RPE cells **(Fig 4A-B, Appendix Fig S3B),** which was further validated through elevated mRNA expression of the pro-apoptotic markers *BAX*, *PUMA* and *NOXA* and the p53 target genes *CDKN1A* and *RRM2B* using RT-qPCR, as well as increased levels of cPARP by immunoblotting (**Fig 4C, D, Appendix Fig S3C-D**). Moreover, the TAS1553-SRA737 combination treatment resulted in reduced pCHK1^S296^, total CHK1, total RPA and RRM2 levels, together with concomitant upregulation of pCHK1^S345^, pRPA32^S33^ and *γ*H2AX levels reflecting increased RS and DNA damage (**Fig 4D, Appendix Fig S3D**). In contrast to the combination with prexasertib, we do observe a significant depletion of RRM2 protein levels upon exposure to TAS1553 in combination with SRA737. In line with previous findings, this can be related to a decrease in p4EBP1 levels upon RRM2/CHK1 inhibition (Goss *et al*, 2021), indicating 4E-BP1 activation, concomitant with a reduction of total RPA, CHK1 and RRM2 protein levels, in parallel to their pharmacological inhibition. This effect may be more pronounced with SRA737 given its higher selectivity for CHK1 compared to prexasertib. Both SRA737 low dose single agent treatment and its combination with low dose TAS1553 effectively reduced CHK1 auto-activation, as indicated by downregulated levels of pCHK1^S296^ compared to TAS1553 (low dose) single or control (DMSO) treatment.

**Figure 4:**
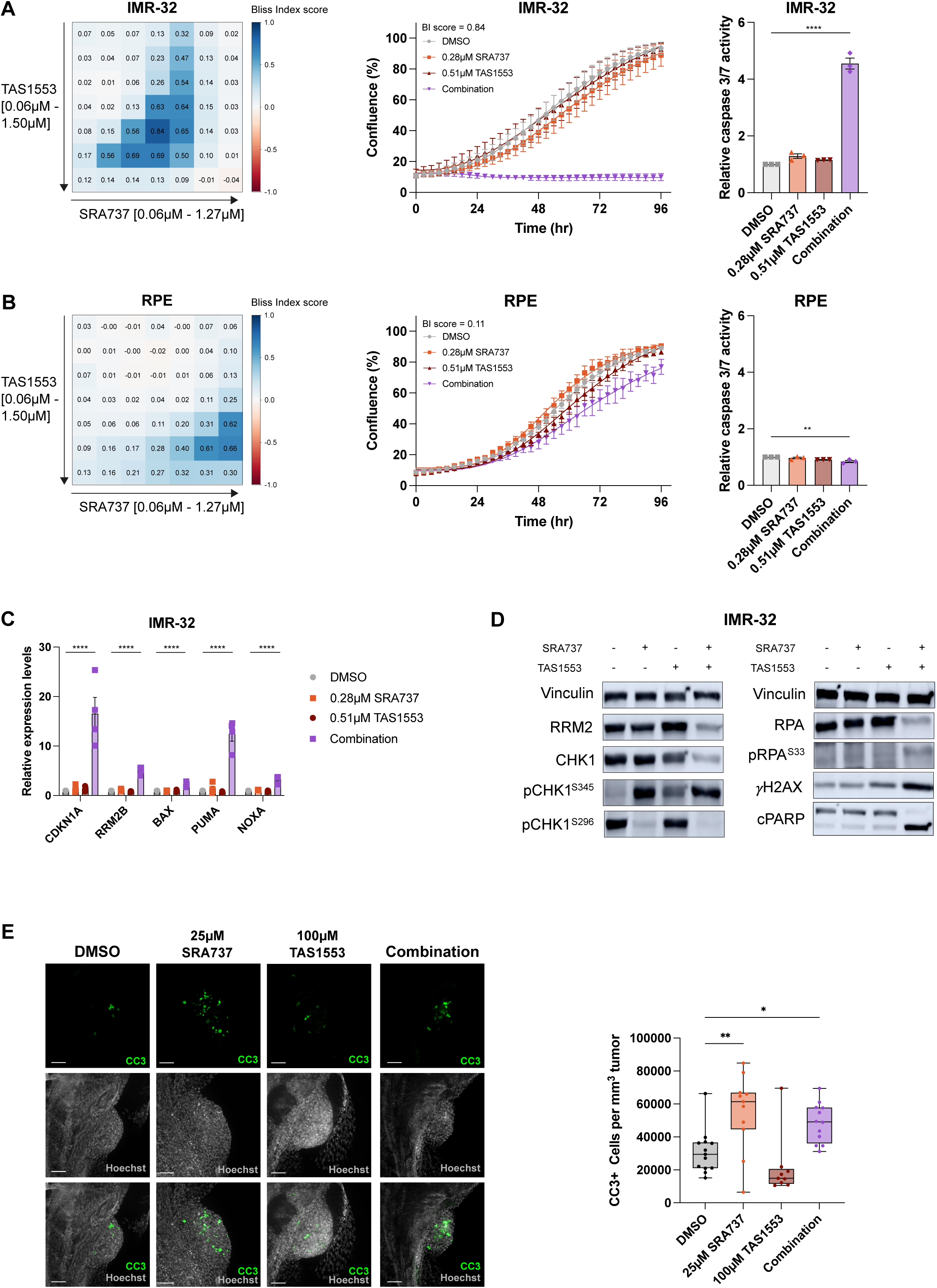
Scrutinizing selective CHK1 inhibition using SRA737 as novel synergistic entry point with TAS1553. A. Heatmap showing the bliss index (BI) scores for IMR-32 after 96h of treatment with a range of concentrations of TAS1553 and SRA737. Biological replicates= 5 in duplicate (*left*). Cell confluency was monitored for 96h and cells were treated with the concentrations with the highest BI score (0.28µM SRA737 and 0.51µM TAS1553). Biological replicates= 5 in duplicate, error bars represent SEM (*middle*). Caspase-Glo luminescence-based measurements of caspase 3/7 activity in IMR-32 following treatment with TAS1553 and SRA737 for 48h. Biological replicates= 3 in triplicate, error bars represent SEM. Significance was measured using the one-way ANOVA followed by a Dunnett’s multiple comparison test (****= p≤0.0001) (*right*). B. Heatmap showing the BI scores for RPE after 96h of treatment with a range of concentrations of TAS1553 and SRA737. Biological replicates= 3 in duplicate (*left*). Cell confluency was monitored for 96h and cells were treated with the concentrations with the highest BI score in NB (0.28µM SRA737 and 0.51µM TAS1553). Biological replicates= 3 in duplicate, error bars represent SEM (*middle*). Caspase-Glo luminescence-based measurements of caspase 3/7 activity in RPE following treatment with TAS1553 and SRA737 for 48h. Biological replicates= 3 in triplicate, error bars represent SEM. Significance was measured using the one-way ANOVA followed by a Dunnett’s multiple comparison test (**= p≤0.01) (*right*). C. RT-qPCR based measurements of mRNA expression of the apoptotic genes *BAX, PUMA* and *NOXA*, and the *p21* target genes *CDKN1A* and *RRM2B* after treatment of IMR-32 with the best combination of TAS1553 (0.51µM) and SRA737 (0.28µM) that resulted in the highest synergy scores at 48h. Biological replicates= 4 in duplicate. Significance was measured using the one-way ANOVA followed by a Dunnett’s multiple comparison test (*= p≤0.05, **= p≤0.01, ***= p≤0.001, ****= p≤0.0001), error bars represent SEM. D. Representative immunoblotting data for various replicative stress and DNA damage markers in IMR-32 following 48h treatment with 0.28µM SRA737 and 0.51µM TAS1553 for 48h. The quantification of all replicates is shown in Supplementary Figure S5B. E. Representative image of zebrafish treated with DMSO or 25µM SRA737 and 100µM TAS1553. CC3 is shown in green and Hoechst in grey, scale bar represents 50µm (*left*). Boxplot showing increased CC3 expression in SK-N-AS NB cells injected in zebrafish after 96h of treatment with 25µM SRA737 and 100µM TAS1553 (*right*).

Next, zebrafish xenografts were treated with the MNTD of the TAS1553-SRA737 combination (100 µM and 25 µM) (**Appendix Fig S5B),** 24 hours after engraftment for 96 hours (treatment was refreshed every 24 hours). Again, we observed elevated levels of CC3+ cells in the zebrafish treated with the combination compared to control and single agent treated zebrafish (**Fig 4E**). Collectively, these results indicate that TAS1553 and SRA737 are potent next generation inhibitors that open a novel avenue for combined selective RRM2-CHK1 inhibition in neuroblastoma and pediatric sarcomas.

### Combined RRM2-CHK1 inhibition leads to differential expression of MYC(N) and ALK controlled gene signatures

In a next step, we scrutinized the molecular consequences imposed by either TAS1553-prexasertib or TAS1553-SRA737 drugging, showing a clear and robust transcriptional effect of the combination treated samples compared to single treatment and DMSO treatment (**Appendix Fig S7A, B**). As anticipated, genes involved in cell death (*CDKN1A, BBC3, BAX*) and DNA damage (*PLK2, GADD45A*) were amongst the top-ranked upregulated genes in both IMR-32 and CLB-GA. Furthermore, the expression of *MDM2*, is upregulated after treatment, which might result from counteracting the reduction in *p53* pathway activity (**Fig 5A, B**) (Yao *et al*, 2024). Similarly to our previously published transcriptome data (Nunes *et al*, 2022), GSEA pointed towards significant downregulation of G_2_-M checkpoint genes, E2F targets and MYC targets while p53 pathway signature genes are upregulated (**Fig 5C**). Moreover, we identified a strong overlap in transcriptional profiles between the different combinations of RRM2 and CHK1 inhibitors evaluated in this study (**Fig 5D, Appendix Fig S7C**). Additionally, we performed GSEA using our different RRM2-CHK1 combination datasets by means of a custom compiled set of publicly available gene signatures (**Fig 5E**). This analysis confirmed significant downregulation of MYCN driven targets in all drug combinations included in our study (Valentijn *et al*., 2012), together with the pan-cancer prexasertib response signature (Blosser *et al*, 2020) and a gene signature associated with elimusertib mediated ATR inhibition in NB (Szydzik *et al*, 2021). Interestingly, genes activated by ALK signaling are upregulated (Lambertz *et al*, 2015). These findings suggest that the RRM2-CHK1 inhibition treatment could be further potentiated with pharmacological ATR or ALK inhibition which should be further evaluated. Finally, we could show that adrenergic markers were downregulated upon combined RRM2-CHK1 inhibition (van Groningen et al, 2017) (**Fig 5E**).

**Figure 5:**
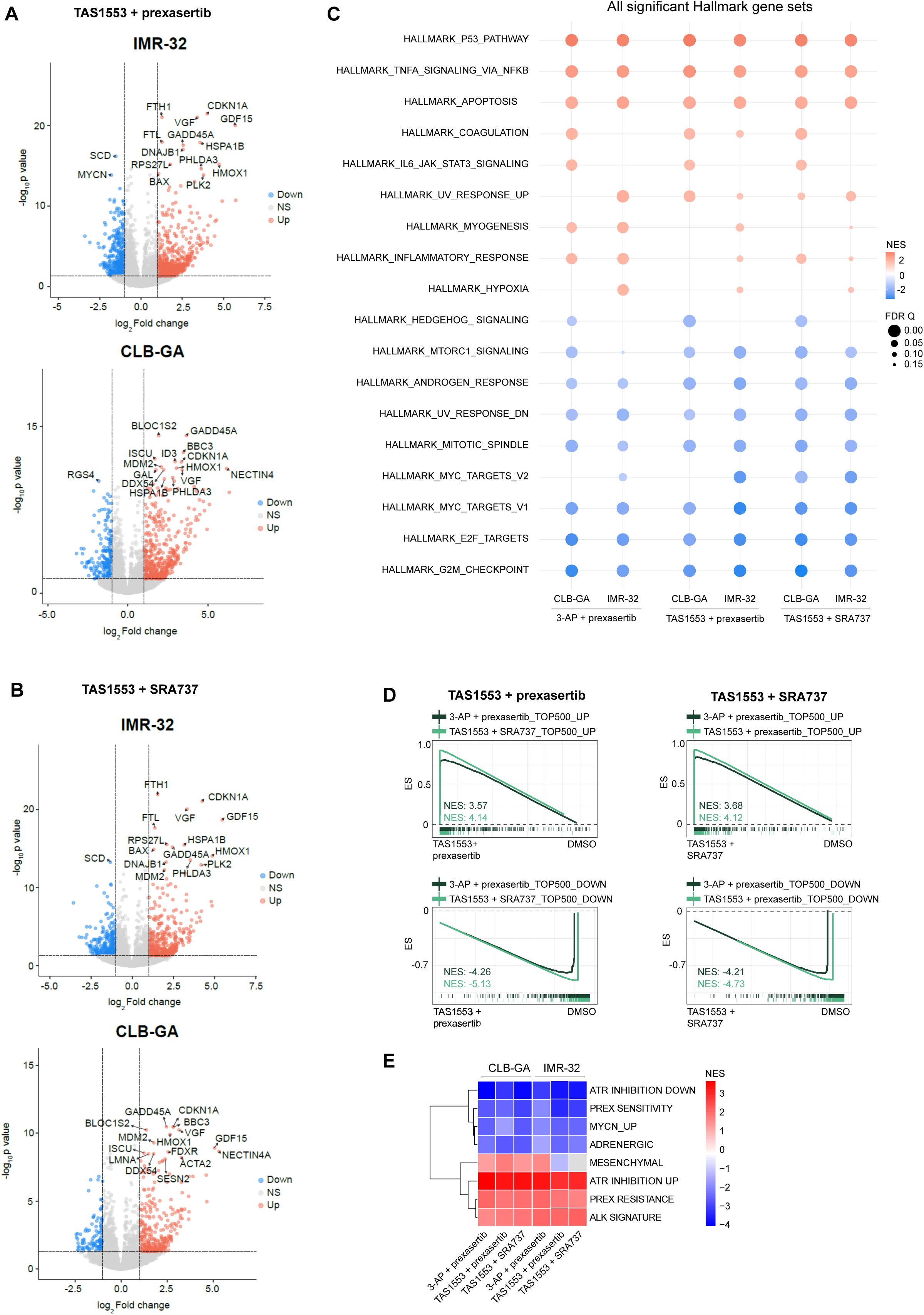
Benchmarking gene expression signatures following TAS1553 exposure in combination with either SRA737 or prexasertib. A. Volcano plots of IMR-32 (*top*) and CLB-GA (*bottom*) of significantly DEGs after treatment with 0.74nM prexasertib and 0.88µM TAS1553 for 48h. Genes indicated by name cover the top 15 DEGs. B. Volcano plots of IMR-32 (*top*) and CLB-GA (*bottom*) of significantly DEGs after treatment with 0.28µM SRA737 and 0.51µM TAS1553 for 48h. Genes indicated by name cover the top 15 DEGs. C. Bubble plot showing the top gene sets enriched after the indicated combination treatment for 48h for IMR-32 and CLB-GA. D. GSEA of the RNA-seq based transcriptome profiles following combined drug treatment with TAS1553 and prexasertib (*left*) or TAS1553 and SRA737 (*right*) in IMR-32 cells showing a significant overlap between genes enriched upon combination treatment of TAS1553 with prexasertib, TAS1553 with SRA737 or 3-AP with prexasertib. E. Heatmap showing the NES scores from our different combination treatments compared to publicly available gene sets (Valentijn *et al*, 2012; Van Groningen *et al*, 2017; Szydzik *et al*, 2021; Blosser *et al*, 2020; Lambertz *et al*, 2015).

### TAS1553 treatment downregulates expression of key RNA splicing components

While the majority of gene signatures associated with the molecular response to RRM2-CHK1 inhibitors included in this study show a significant overlap (**Fig 2B** and **Fig 5C**), we observed with both TAS1553 single treatment and in combination with CHK1 inhibition a more pronounced downregulated expression of genes encoding core spliceosome components (**Fig 6A, Appendix Fig S8A**), including the small nuclear ribonucleoprotein (snRNP) U1 and U5 (**Fig 6B**). While U1 snRNP recognizes the 5’ splicing site (5’ SS) and is necessary to initiate the splicing process via the spliceosome (Wilkinson *et al*, 2020), U5 snRNP interacts with U4/U6 to form the complete spliceosome complex and start the branching and exon ligation steps (Nguyen *et al*, 2015). Furthermore, *Prp43*, involved in the disassembly of the spliceosome, is also downregulated suggesting a depletion of U2/U5/U6 to form new spliceosomes (**Fig 6B**). Indeed, previous studies have indicated a functional interconnection between pre-mRNA processing and RS (Shkreta & Chabot, 2015; Paulsen *et al*, 2009). It can therefore be anticipated that TAS1553 could also act synergistically with splicing inhibitors and should be further investigated in follow-up of this study.

**Figure 6:**
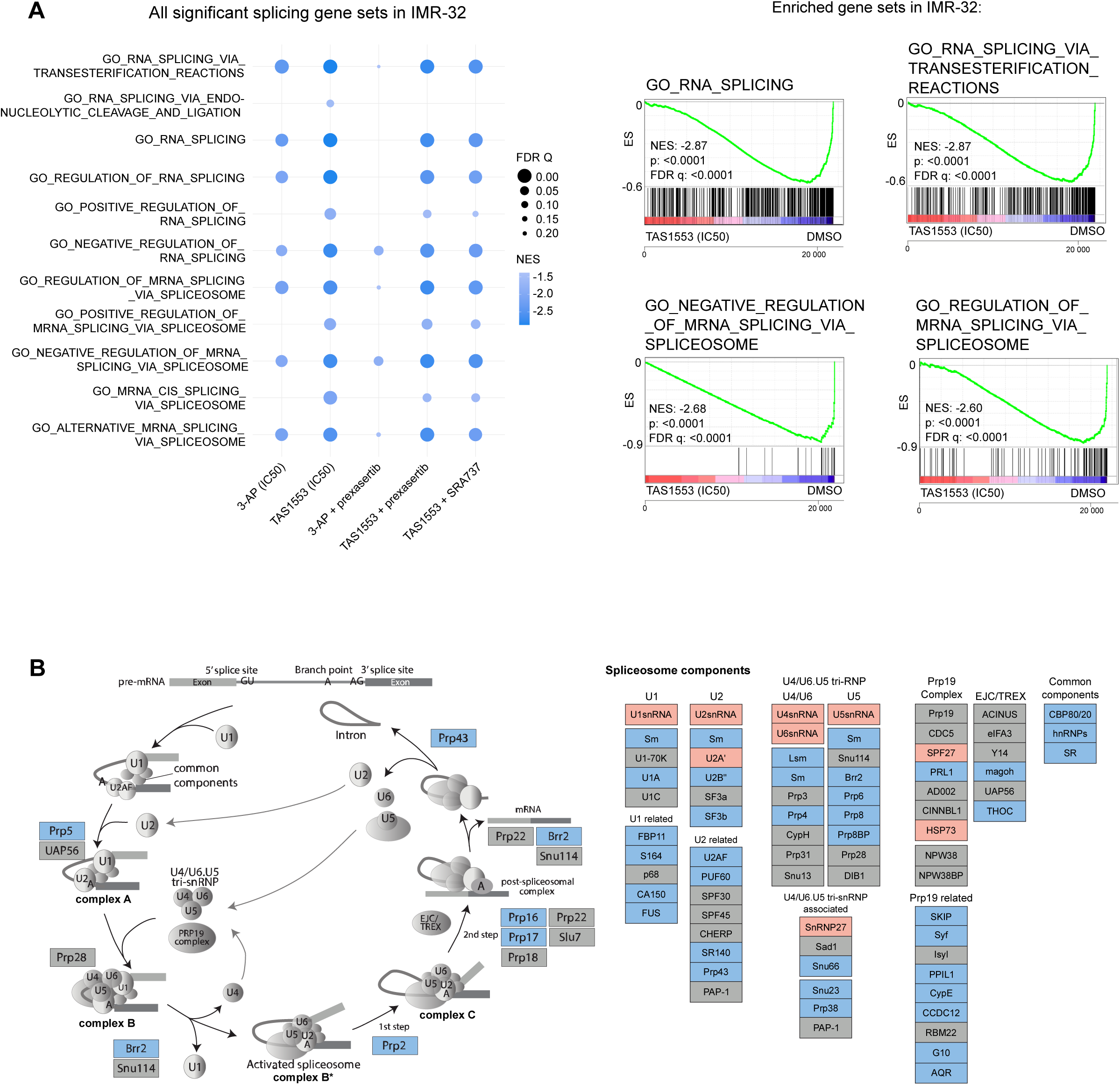
TAS1553 induces downregulation of genes encoding splicing pathway components. A. Bubble plot showing significant downregulation of gene sets involved in RNA splicing upon treatment with TAS1553 or the combination with CHK1 inhibitors in IMR-32 (*left*). Corresponding GSEA of two gene sets involved in RNA splicing (top) or the spliceosome (bottom) after treatment with IC_50_ of TAS1553 in IMR-32 (*right*). B. KEGG pathway analysis of the spliceosome in IMR-32 indicating which genes are significantly differentially expressed (*blue:* downregulated by TAS1553; *red:* upregulated by TAS1553) upon treatment with the IC_50_ of TAS1553 for 48h.

## Discussion

We evaluated the *in vitro* and *in vivo* phenotypic consequences of selective RNR inhibition through disruption of the RRM1-RRM2 complex using the novel TAS1553 inhibitor (Ueno *et al*, 2022). At the molecular level, TAS1553 treatment induced elevated levels of RS and DNA damage and showed synergism with the CHK1 inhibitors prexasertib, in line with our previous findings (Nunes et al, 2022) and now also SRA737. Interestingly, transcriptome analysis revealed that TAS1553 either as single agent or in combination with CHK1 inhibition resulted in selective downregulation of gene sets related to RNA splicing components.

We and others previously reported on the dependency role of RRM2 in tumorigenesis, establishing it as an attractive therapeutic target for extensive drug development efforts (Bokhari *et al*, 2022; Ohmura *et al*, 2021; Mazzu *et al*, 2019; Nunes *et al*, 2022). First generation RRM2 inhibitors included hydroxy-urea (HU) and 3-AP. In short, HU acts as radical scavenger and was approved by the Food and Drug Administration (FDA) for treatment of sickle cell disease and myeloid neoplasms (Fishbein *et al*, 1964; Hankins *et al*, 2014). However, HU has a short half-life in humans and tumor cells acquire resistance to HU via increased expression of RRM2 (Yen *et al*, 1994; Zhou *et al*, 1998). In contrast, 3-AP is an iron chelator (Atkin et al, 1973; Shao et al, 2006) and and able to overcome this resistance (Finch *et al*, 2000), but further clinical development efforts have been halted. In contrast, the RRM1 inhibitor gemcitabine is a nucleoside analog for dCTP and can bind to the catalytic site of RRM1 (Plunkett *et al*, 1991; Heinemann *et al*, 1990; Xu *et al*, 2006). However, RRM1 is expressed throughout the cell cycle, while RRM2 expression is induced in early S-phase and rapidly degraded upon S-G2 phase transition (Engström *et al*, 1985). Therefore, RRM2 is the rate limiting factor of the RNR complex and considered the most clinically relevant RNR target protein. Next-generation and more on-target RRM2 inhibitors were considered to be required to overcome therapy resistance and decrease the number of adverse events. In this respect, Ueno and colleagues developed TAS1553 which specifically disrupts the interaction between RRM1 and RRM2 (Ueno et al, 2022).

In addition, tumor cells rapidly acquire resistance against single compound treatment regimens (Mokhtari *et al*, 2017). Therefore, we aimed in this study to evaluate both TAS1553 as single agent and in combination with the CHK1 inhibitors prexasertib and SRA737, the latter two both being ATP-competitive CHK1 protein kinase inhibitors. Notably, synergistic interactions with SRA737 could be achieved with a lower input concentration of TAS1553 compared to the combination with prexasertib. This could be due to the stronger selectivity of SRA737 for CHK1 compared to prexasertib (1,000-fold greater selectivity) (King *et al*, 2015; Osborne *et al*, 2016). However, this selectivity profile was determined by *in vitro* kinase assays. Interestingly, a study by Ditano *et al*. compared the CHK1 versus CHK2 selectivity of prexasertib versus SRA737 and suggests that the prexasertib concentrations used in our study do not affect CHK2, therefore implying that presumed selectivity for CHK1 over CHK2 of SRA737 does not explain the observed stronger synergistic interaction of TAS1553 with SRA737 versus prexasertib.

Pharmacological targeting of the RS and DNA damage response pathway results in fork collapse and double stranded DNA (dsDNA) breaks, resulting in release of dsDNA into the cytosol, which can lead to activation of cGAS-STING pathway as previous studies shown using poly(ADP-ribose) polymerase (PARP) inhibitors (Schoonen *et al*, 2019; Chabanon *et al*, 2019). More recently, the study of Taniguchi *et al*. showed activation of cGAS-STING following ATR inhibition in small-cell lung cancer via DNA damage mediated micronuclei (Taniguchi *et al*, 2024). While NB tumors are classified as immunologically ‘cold’ tumors (Blavier *et al*, 2020), we showed that combined RRM2-CHK1 inhibition resulted in a significant upregulation of genes involved in inflammatory and interferon alpha responses. Therefore, this could be a promising entry point to sensitize NB tumors for immune checkpoint inhibitors.

Notably, we also identified that TAS1553 exposure unique imposes a significant downregulation of RNA splicing and previous studies indeed support functional interaction between genomic stability and pre-mRNA processing. In this respect, it has been shown that DNA damage signaling, amongst others via ATR, can regulate splicing events, via perturbed acetylation or phosphorylation of splicing factors (Shkreta & Chabot, 2015). On the other hand, downregulation of genes involved in mRNA processing results in increased levels of *γ*H2AX (Paulsen *et al*, 2009). Here, treatment with TAS1553 lead to selective downregulation of genes necessary to form small nuclear ribonucleoproteins (snRNP; U1, U2, U4, U5 and U6). Previous studies have indeed shown that mutated U2 snRNP results in an increase of numbers of R-loops (Tanikawa *et al*, 2016; Nguyen *et al*, 2018) and that depletion of the U2 snRNP leads to decreased expression of the DDR factors RAD51, ATM and CHK1 (Tanikawa *et al*, 2016). Interestingly, RNAi screening data of the DepMap initiative (Broad Institute, depmap.org) point towards SF3B1 as selective NB dependency compared to other solid pediatric tumor entities (**Appendix Fig S8B**). Therefore, we anticipate that pharmacological SF3B1 inhibition could act synergistically with TAS1553 and should be further investigated in follow-up of this study.

In conclusion, we show that the RNR disruptive agent TAS1553 is a potent novel therapeutic entry point for NB and pediatric sarcoma cell lines and *in vivo* in a NB zebrafish xenograft model as single agent and in combination with CHK1 inhibition. Future studies should consider to exploit combined RRM2-CHK1 for immunomodulation in NB and recommend TAS1553 as potent entry point for novel synergistic drug interactions with splicing inhibitors for pediatric solid tumor entities.

## Materials and Methods

### Cell culture

The human NB cell lines CLB-GA, SK-N-AS, SH-SY5Y, SH-EP, CHP-212, IMR-32, N206, NGP, SK-N-BE(2)-C and TR14 were cultured in RPMI1640 (Gibco #52400041), supplemented with 10% fetal bovine serum (FBS) (Sigma-Aldrich #F0804), 2mM L-glutamine (Gibco #11548876) and 100IU/mL penicillin/streptomycin (Gibco #11548876). RPE cells were cultured in DMEM-F12 (Gibco #11594426), supplemented with 10% FBS, 2mM L-glutamine and 100IU/mL penicillin/streptomycin. Human Ewing sarcoma (EWS) cell lines A-673, ES7, EW-7, MHH-ES-1, TC-71, human osteosarcoma cell lines CAL-72, HS-Os-1, HuO9, MG-63, SaOS-2, U2OS were maintained in RPMI1640 medium with stable glutamine (Sigma-Aldrich #R8758) supplemented with 10% tetracycline-free FBS (Sigma-Aldrich #F7524), 100 IU/ml penicillin and 100 µg/ml streptomycin (Sigma-Aldrich #P4333). The human RMS cell lines Rh41 and RMS-YM were cultured in RPMI1640 (Gibco #52400041), supplemented with 10% FBS (Tico #FBSEU500), 2mM L-glutamine (Gibco #11548876) and 100IU/mL penicillin/streptomycin (Gibco #11548876). The human RMS cell lines RD, Rh30 and RMS-1 were cultured in DMEM (Gibco #41965-039), supplemented with 10% FBS (Tico #FBSEU500), 0.1mM non-essential amino acids (NEAA) (Gibco #12084947), 2mM L-glutamine (Gibco #11548876) and 100IU/mL penicillin/streptomycin (Gibco #11548876). All cells were incubated at 37°C and 5% CO_2_. Cell line characteristics can be found in supplementary tables S1, S2, S3 and S4.

### Compounds

TAS1553 (#HY-150785) and prexasertib (#HY-18174A) were purchased via Bio-Connect. SRA737 (#S8253) was purchased via Selleckchem.

### Cell confluency assay

NB cell lines were seeded in 384-well plates at a density ranging from 1,500-6,000 cells/well and allowed to adhere overnight. RPE cells were seeded at a density of 200 cells/well. RMS cell lines were seeded at a density of 1,000-3,000 cells/well. Compounds were added in a concentration range to the plates using the Tecan D300e Digital Dispenser. Each experiment was performed with three technical replicates and at least three biological replicates were collected. The cells were monitored over time in the IncuCyte Live Cell Imaging System for at least 96h. Each image was analyzed through the IncuCyte Software and cell proliferation was determined by the area (percentage of confluence) of cell images over time. Cell confluency was normalized to the DMSO control at 72h or 96h and the IC_50_ was determined using HTSplotter (Nunes *et al*, 2024). Graphs were made using GraphPad (version 10.3.1) and dose-response curves were determined using the variable slope (four parameters) equation. The error bars indicate standard error of the mean (SEM) of at least three biological replicates.

### Cell viability assay

EWS or OS cell lines were seeded in a 96-well plate at a density ranging from 1,000-5,000 cells/well and allowed to adhere overnight. Compounds were serially diluted and added to the plates in a total volume of 100µL with at least three technical replicates. Cell viability was assessed 72h after the treatment start with Resazurin cell viability assays (16µg/mL, Sigma-Aldrich). Dose response curves were simulated by nonlinear regression models using the variable slope equation and IC_50_ values were calculated using GraphPad (version 10.3.1) by normalizing to the respective controls (vehicle alone).

### dNTP analysis

IMR-32 cells were seeded at a density of 3×10^5^ cells in a 6-well plate and allowed to adhere overnight. The following day, medium was refreshed with medium supplemented with the compounds or medium only and again incubated for 48h. Before harvesting, the medium was collected, plates were put on ice and cells were washed with 1mL fresh prepared ice-cold 0.9% NaCl. Next, extraction buffer was added and incubated on ice for 3 min. This was followed by scraping the cells and samples were stored overnight at -

80°C. The next day, samples were centrifuged for 5 min, 20,000g at 4°C. The supernatant was collected and sent for dNTP analysis. The cell pellets were stored at -80°C to determine the protein concentrations. Cell pellets were lysed in 200µL 200mM NaOH and heated to 95°C for 20 min. Next, samples were centrifuged for 10min, 6000g at 4°C and the supernatant was collected. Protein concentrations were determined using the Pierce^TM^ BCA Protein Assay Kit (ThermoFisher #23227).

### Combination and synergism measurements

CLB-GA and IMR-32 were seeded in 384-well plates at a density ranging from 1,500-2,000 cells/well and allowed to adhere overnight. The RMS cell lines Rh30 and RM-SYM were seeded at a density ranging from 1,500-3,000 cells/well and allowed to adhere overnight. The following day, cells were treated with the respective compounds alone or in a combination matrix using the Tecan D300e Digital Dispenser. Each experiment was performed with two technical replicates. The cells were monitored over time in the IncuCyte Live Cell Imaging System for at least 96h. Each image was analyzed through the IncuCyte Software and cell proliferation was determined by the area (percentage of confluence) of cell images over time. Synergism was determined using the Bliss Index (BI) score as identified by HTSplotter (Nunes *et al*, 2024). Each experiment was repeated 3-5 times to determine combination with the highest BI score. Confluency graphs were made using GraphPad (version 10.3.1). The error bars indicate SEM.

For EWS, A-673 and ES7, cells were seeded in a 96-well plate in triplicate at a density of 2,500 cells/well and 1,500 cells/well, respectively, and allowed to adhere overnight. The following day, cells were treated with the indicated drug combinations in three serial doses (4×4 matrices). Cell viability was assessed with Resazurin cell viability assays 72h after the treatment start and values were normalized to the respective controls (vehicle alone).

For OS, MG-63 and U2OS cells were seeded in a 96-well plate in duplicate at a density of 1,000 cells/well and 1,500 cells/well, respectively, and allowed to adhere overnight. The following day, cells were treated with various drug combinations in five serial doses (6×6 matrices). Cell viability was assessed with Resazurin cell viability assays 72h after the treatment start and values were normalized to the respective controls (vehicle alone). Synergism was determined using the BI score as identified by HTSplotter (Nunes *et al*, 2024).

### Apoptosis assay

CLB-GA (15,000 cells/well), IMR-32 (14,000 cells/well) and RPE (2,000 cells/well) were seeded in a 96-well plate and allowed to adhere overnight. The next day, cells were treated with the designated compounds using the Tecan D300e Digital Dispenser and put in the incubator. Pictures were taken 0h, 24h and 48h after treatment with the IncuCyte Live Cell Imaging System to follow cell growth. After 48h, apoptosis levels were measured using the Caspase-Glo 3/7® Assay (Promega #G8091). The manufacturer’s protocol was adapted, by adding 50µL of reagent per well. Each experiment included three technical replicates. The average luminescence of the treated wells was scaled to the DMSO control. Graphs were made using GraphPad (version 10.3.1) and error bars represent the SEM of three biological replicates. Significance was determined using the one-way ANOVA followed by a Dunnett’s multiple comparison test.

### RNA isolation and RT-qPCR

IMR-32 and CLB-GA cells were seeded in T25 culture flasks (Greiner #690175) at a density of 0.8×10^6^ and 1×10^6^ cells respectively and allowed to adhere for 48h. After 48h, medium was refreshed with medium supplemented with DMSO, the compounds or medium only and again incubated for 48h. Next, cells were harvested using a cell scraper and ice-cold phosphate buffered saline (PBS, Gibco #14190094), and centrifuged for 5 min, 2500 rpm, at 4°C. This was followed by a wash in PBS and again centrifuged for 5 min, 2500 rpm, at 4°C. Cell pellets were dissolved in QIAzol Lysis Reagent (Qiagen #79306) and snap frozen in liquid nitrogen.

RNA isolation was performed using the miRNeasy mini kit (Qiagen #217004) using the manufacturer’s protocol, including the on-column DNase treatment (Qiagen #79254). Concentrations were measured using the NanoDrop (Thermo Fisher Scientific). Complementary DNA (cDNA) was made using the iScript Advanced cDNA synthesis kit (Bio-Rad #172-5038). The PCR mastermix was composed out of 5 ng cDNA, 2.5µL SsoAdvanced Universal SYBR Green Supermix (Bio-Rad #172-5274) and 0.25µL of forward and reverse primers (sequences can be found in supplementary table S6). RT-qPCR was performed using the LC-480 device (Roche). The gene expression analysis was performed using Qbase+ (CellCarta, version 3.4) and the average gene expression was scaled to the DMSO control. Graphs were made using GraphPad (version 10.3.1), error bars represent the SEM of four biological replicates and statistical analysis was performed using the multiple unpaired T-test for the single treated experiments and one-way ANOVA followed by a Dunnett’s multiple comparison test for the combination treatment.

### Protein isolation

The NB cell lines IMR-32 and CLB-GA were seeded and drugged as described in the ‘RNA isolation and RT-qPCR’ section. However, cell pellets were snap frozen in liquid nitrogen after centrifugation.

Cell pellets were lysed in cold RIPA buffer (250 mg sodium deoxycholate, 150mM NaCl, 50mM Tris-HCl pH 7.5, 0.01% SDS solution, 0.1% NP-40), supplemented with protease (Roche #11836145001) and phosphatase (Roche #04906845001) inhibitors. This was followed by rotation at 4°C for 1h to achieve more complete lysis. Next, samples were centrifuged for 10 min, 10,000 rpm, at 4°C, and the lysates were collected. Protein concentrations were determined using the Pierce^TM^ BCA Protein Assay Kit (ThermoFisher #23227).

### Western blotting

Prior to gel loading, 30µg of protein was denaturated by adding 5x Laemmli denaturation buffer supplemented with β-mercaptoethanol, and heated to 95°C for 10min. Next, samples were loaded on 10% SDS-PAGE gels with 10x Tris/glycine/SDS buffer (Bio-Rad #1610772) and run for 1h at 130V. This was followed by blotting on nitrocellulose membranes in 10% of 10x Tris/Glycine buffer (Bio-Rad #1610771) and 20% methanol, for 1h at 100V. Ponceau staining was used to confirm the protein transfer. Membranes were blocked in 5% milk or 5% BSA (Sigma-Aldrich #A7030) in TBS-T for 1h at room temperature (RT). Primary antibody dilutions were done in blocking buffer overnight at 4°C. The next day, blots were washed 3x for 5min with TBS-T followed by 1h incubation of the secondary antibodies. Subsequently, blots were washed again 3x for 5min with TBS-T. The immunoblots were visualized on the Amersham^TM^ Imager 680, using the enhanced chemiluminescent West Femto (Thermo Scientific #34096) or West Dura (Fisher Scientific #10220294). To visualize additional antibodies, blots were stripped using the Restore^TM^ Stripping Buffer (Thermo Scientific #10016433). The antibody information can be found in supplementary table S7. Quantification analysis was performed through ImageJ software (version 2.14.0). Fold changes were scaled to vinculin. Graphs were made using GraphPad (version 10.3.1), error bars represent the standard deviation of three biological replicates and statistical analysis was performed using the unpaired T-test for the single treated experiments and one-way ANOVA followed by a Dunnett’s multiple comparison test for the combination.

### RNA sequencing

RNA was isolated as described in the ‘RNA isolation and RT-qPCR’ section. RNA quality was measured using the Fragment analyzer 12 capillary array prior to profiling. A total of 10ng of RNA was used as input for the library preparation with the QuantSeq-Pool Sample-Barcoded 3’ mRNA-Seq Kit (Lexogen #139.96). The manufacturer’s protocol was adapted to use only half of the total volumes of the solvents, and 15 PCR cycles were used for amplification. qPCR quantification of the libraries was done using the Kapa Library Quantification Kit for the Illumina LightCycler 480 (Roche diagnostics #07960298001). RNA-seq libraries were sequenced on the Illumina NextSeq 2000 platform using the NextSeq 2000 P3 Reagents 100 Cycles paired-end kit (Illumina #20040559). Data analysis followed the workflow outlined on the Lexogen GitHub repository (https://github.com/Lexogen-Tools/quantseqpool_analysis). Briefly, sample and read quality were evaluated with FastQC (v0.11.7). The FASTQ files were aligned to the Homo sapiens reference genome (GRCh38.96) using STAR (v2.7.2b). Post-alignment, read counts were generated and normalized with edgeR (v3.36.0). After performing PCA plotting, certain replicates were removed from the analysis as they did not cluster with the other replicates, indicating potential outliers or inconsistencies.

Differential expression analysis was performed using the Limma-Voom pipeline (v3.50.3) with three or four replicates per condition. Only genes with an adjusted P-value < 0.05 and a logFC of ≤ -1 (downregulated) or ≥ 1 (upregulated) were considered significant. Gene Set Enrichment Analysis (GSEA) was performed on genes ordered according to the differential expression statistic (t) value. GSEA utilized gene sets from MSigDB (https://www.gsea-msigdb.org/gsea/msigdb/) and custom gene lists from this and previous studies (3-AP datasets: GSE161902 and ALK signaling signature (Lambertz *et al*, 2015)). After GSEA, only protein-coding genes were retained for further analysis. All figures were generated using ggplot2 (v3.5.1), pheatmap (v1.0.12) and pathview (v1.42.0) packages in R.

All RNA-seq data are available through the Gene Expression Omnibus (GEO) repository (GSE279147).

### Zebrafish housing

All procedures involving experimental animals were approved by the local animal ethics committee (Ghent University hospital, Ghent, Belgium), permit number ECD 17-100 and performed according to local guidelines and policies in compliance with national and European law.

Zebrafish were maintained in a Zebtec semi-closed recirculation housing system (Techniplast). The water had a constant temperature (27-28 °C), pH (∼7.5), conductivity (∼500μS) and light/dark cycle (14h/10h).

### Determining the maximum tolerable dose (MTD)

Wild type (AB) zebrafish at 3 days post fertilization were randomly distributed into groups and treated by submersion in different concentrations of TAS1553 and the CHK1 inhibitors prexasertib or SRA737 or DMSO diluted in E3 medium. Fish were exposed to treatment for 96h and the E3 medium with compound was refreshed every 24 hours. Every day, the zebrafish were screened for signs of toxicity. All zebrafish were euthanized with tricain 25x (Sigma-Aldrich #A5040-25G) after 96h of treatment. The MTD was defined as the highest concentration of TAS1553 and CHK1 inhibition that did not cause any signs of toxicity, including death, cardiac edema, a slower heart rate and a curved tail.

### Establishment of zebrafish avatars

SK-N-AS cells were grown to 70% confluence in a T75 flask and detached from the surface using trypsin (Gibco #11580626). Next, the cells were fluorescently stained with 4µl/ml Vybrant CM-DiI (ThermoFisher Scientific #C-7001) in PBS for 10min at 37°C and 10min on ice. The cells were then washed with PBS and resuspended to a final concentration of 0.3×10^6^ cells/µl in PBS. The cells were kept on ice during the injections. Thin wall glass capillaries (World precision instruments #TW100-4) were pulled into microinjection needles using a capillary puller (Sutter instrument P-1000). Wild type (AB) zebrafish at 2 days post fertilization were anaesthetized with tricain 1x in E3 medium for 5min and immobilized in a thin layer of low melting point agarose 1% (Life Technologies Europe #16520-05S0) on a cover slip. Next, the fluorescently labelled cells were injected into the perivitelline space of the zebrafish using the pulled needles and a FemtoJet^®^ 4i microinjector (Eppendorf). The zebrafish were removed from the agarose and sedated for 10 more minutes before they were transferred to E3 medium and incubated at 34°C. Zebrafish xenografts were evaluated 24h post-injection with a NikonSMZ18 stereoscope and randomly distributed into two groups. Zebrafish xenografts were treated for 96h with either 0.5% DMSO or 200µM TAS1553, which was the maximum tolerated dose, by adding the compound to the E3 medium. E3 medium with compound was refreshed every 24h. The treatment was finished 5 days post-injection and the zebrafish xenografts were euthanized with tricain 25x.

### Whole mount immunofluorescent staining

Whole mount immunofluorescent staining was performed as described by M. Martinez-Lopez et al. (Martinez-Lopez *et al*, 2021). In short, zebrafish xenografts were fixed with 4% paraformaldehyde (PFA) / 0.1% Triton X-100 at 4°C overnight and subsequently stored in methanol at -20°C. Zebrafish xenografts were then rehydrated by decreasing methanol concentrations, washed with PBS/0.1% Triton X-100 and bleached for 5min with bleaching solution consisting of 30% H_2_O_2_ (Merck Life Science #H1009) and 10% KOH in dH_2_O. They were subsequently permeabilized by 7min incubation in ice cold 100% acetone and blocked for 1h at RT with blocking buffer consisting of 0.01g/ml BSA, 1% DMSO, 0.5% Triton X-100 and 0.015% goat serum diluted in 1x PBS. Next, xenografts were incubated 1h at RT and overnight at 4°C with primary antibody rabbit anti-cleaved caspase-3 (Cell Signaling Technologies #9661S) diluted 1:100 in blocking buffer. The samples were washed the next day with PBS/0.1% Triton X-100 and PBS/0.05% Tween and incubated 1h at room temperature and overnight at 4°C with secondary antibody Alexa Fluor^®^ 488 goat anti-rabbit IgG (Life Technologies #A11034) diluted at 1:400 and 1% Hoechst in blocking buffer. Afterwards, the zebrafish xenografts were washed with PBS/0.05% Tween, fixed with 4% PFA/0.1% Triton X-100 for 20min and mounted between two coverslips with Mowiol (Sigma-Aldrich 81381-50G). Images were taken with a Zeiss Spinning Disk confocal microscope with 10x water objective and 5µM z-stack interval.

### Cleaved caspase-3 quantification

Images were analyzed using FIJI/ImageJ (version 1.53c). First, the tumor volume for each xenograft tumor was obtained by defining and measuring the tumor area of each Z-stack, multiplying it with the 5µM Z-stack interval and adding up these Z-stack volumes. Next, the total number of cleaved caspase-3 positive cells was counted for each tumor using the Cell Counter plug-in of FIJI/ImageJ and this count was subsequently divided by the tumor volume to obtain the number of cleaved caspase-3 positive cells per mm^3^ tumor. The data shown consists of three biological replicates.

## Data availability

The RNA sequencing data obtained during this study are available through GEO repository with the accession number GSE279147 (https://www.ncbi.nlm.nih.gov/geo/query/acc.cgi?acc=GSE279147).

## Acknowledgments

We would like to thank dr. Igor Fijalkowski and prof. dr. Panagiotis Ntziachristos for the guidance and support with their expertise in RNA splicing. Furthermore, we thank the Zebrafish Core Facility for support and care taking of the zebrafish.

This work was supported by Villa Joep (STI.DIV.2020.0019.01); Kom op Tegen Kanker (KOTK) (STI.VLK.2022.0003.01); Hubert Gouin (INT.DIV.2023.0031.01); Kinderkanker Fonds (KKF) (STI.DIV.2023.0013.01); Stichting Tegen Kanker (STK) (STI.STK.2023.0017.01) and Olivia Fund. The laboratory of T.G.P.G. acknowledges funding of the Barbara and Wilfried Mohr foundation as well as of the European Union (ERC, CANCER-HARAKIRI, 101122595). Views and opinions expressed are however those of the authors only and do not necessarily reflect those of the European Union or the European Research Council.

## Author contributions

Conceptualization/Methodology: I.N., S.L., S.O., T.G.P.G., F.S., A.V.H., L.D., K.D.

Investigation: I.N., S.L., F.D.V., F.M., A.P.E.R., L.V.

Analysis: I.N., S.L., S.B., S.O., L.D.

Visualization: I.N., S.L., S.B., L.D.

Writing – Original draft: I.N., S.L., S.B., S.O.

Writing – Review & Editing: I.N., S.L., S.O., T.G.P.G., F.S., N.V.R., B.D.W., F.S., A.V.H., L.D., K.D.

Funding: T.G.P.G., F.S., K.D.

---

**Figure S1:**
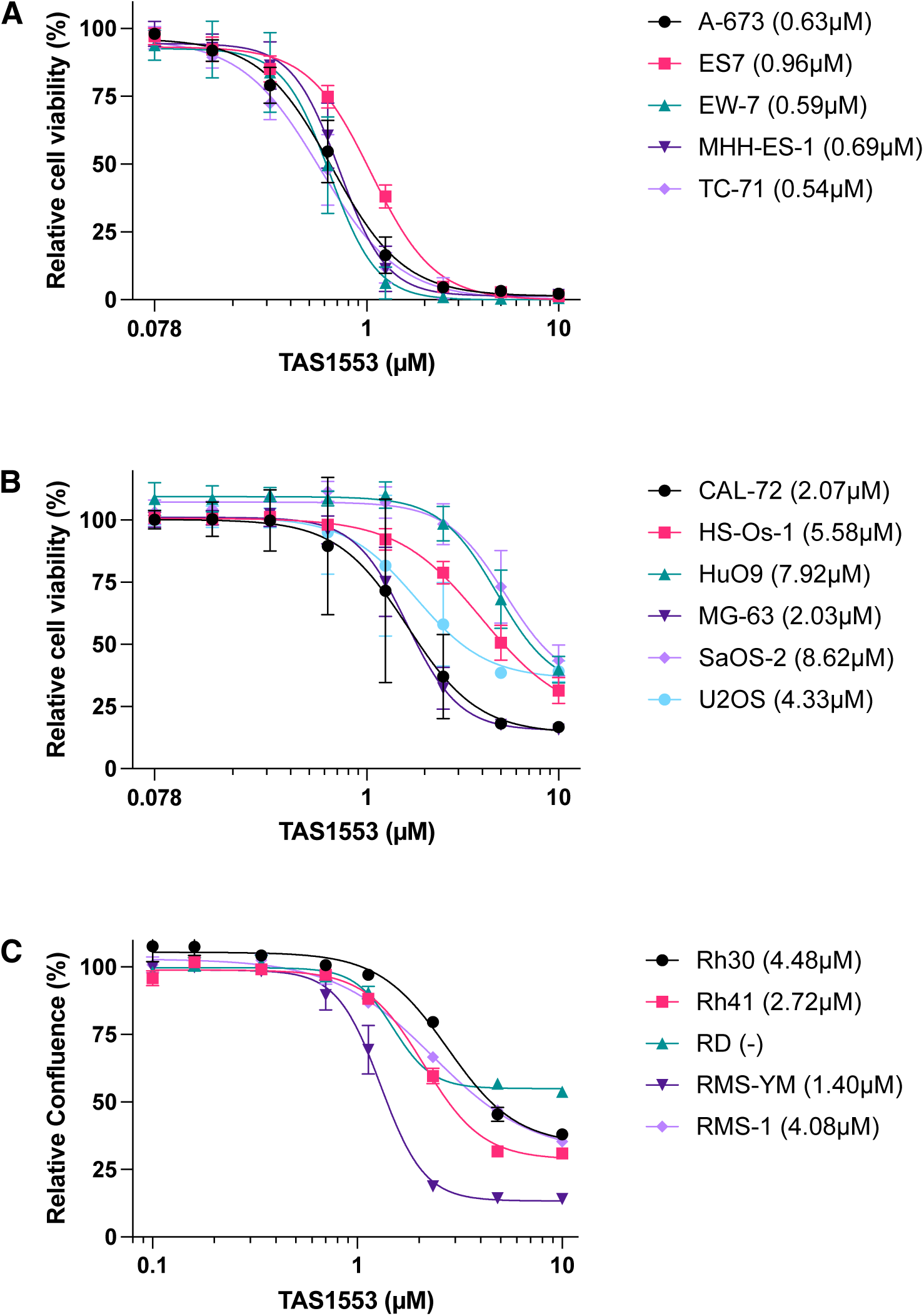
TAS1553 reduces pediatric sarcoma cell growth *in vitro*. A. Dose-response curves of 5 Ewing sarcoma cell lines and their corresponding IC_50_ concentration at 72h. Biological replicates= 4 in triplicate, error bars represent standard error of the mean (SEM). B. Dose-response curves of 6 osteosarcoma cell lines and their corresponding IC_50_ concentration at 72h. Biological replicates= 4 in triplicate, error bars represent SEM. C. Dose-response curves of 5 rhabdomyosarcoma cell lines and their respective IC_50_ concentrations at 72h. Graphs represent the mean of 3-4 biological replicates in triplicate ± SEM.

**Figure S2:**
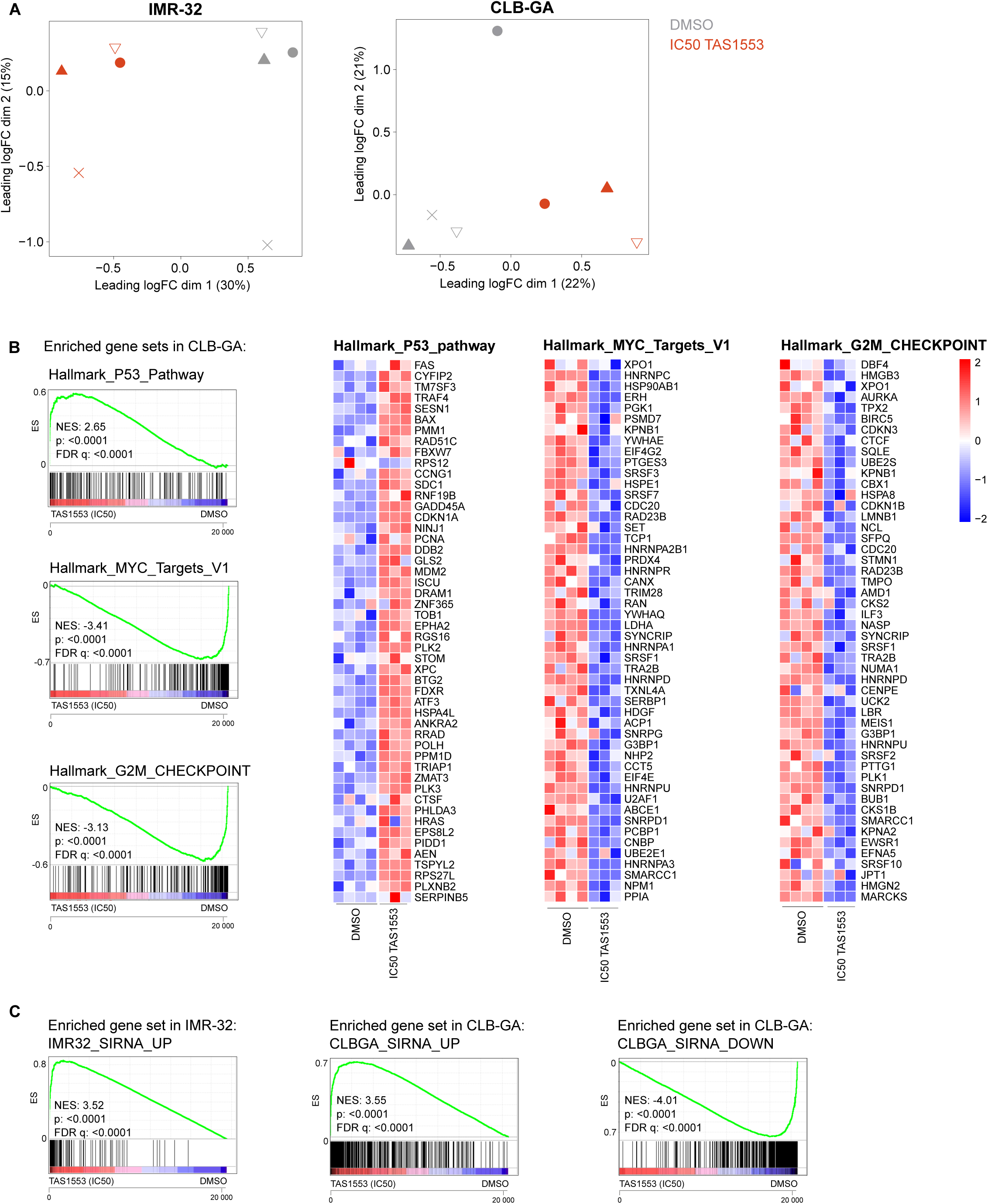
Hallmark gene set enrichment in CLB-GA neuroblastoma cells following TAS1553 treatment. A. Principle component analysis (PCA) showing a clear distinction between cells treated with the control (DMSO) and the IC_50_ of TAS1553 for 48hr for IMR-32 (*left*) and CLB-GA (*right*) (1.18µM and 1.54µM respectively). Symbols represent different replicates. B. GSEA of the RNA-seq based transcriptome profiles after treatment of CLB-GA with 1.54µM TAS1553 (IC_50_) treatment using the Hallmark MSigDB gene sets (*left*) and heatmaps showing the top 50 genes within the core enrichment (*right*). ES = enrichment score. C. GSEA shows an overlap in genes enriched upon siRRM2 treatment and cells treated with the IC_50_ of TAS1553. ES = enrichment score.

**Figure S3:**
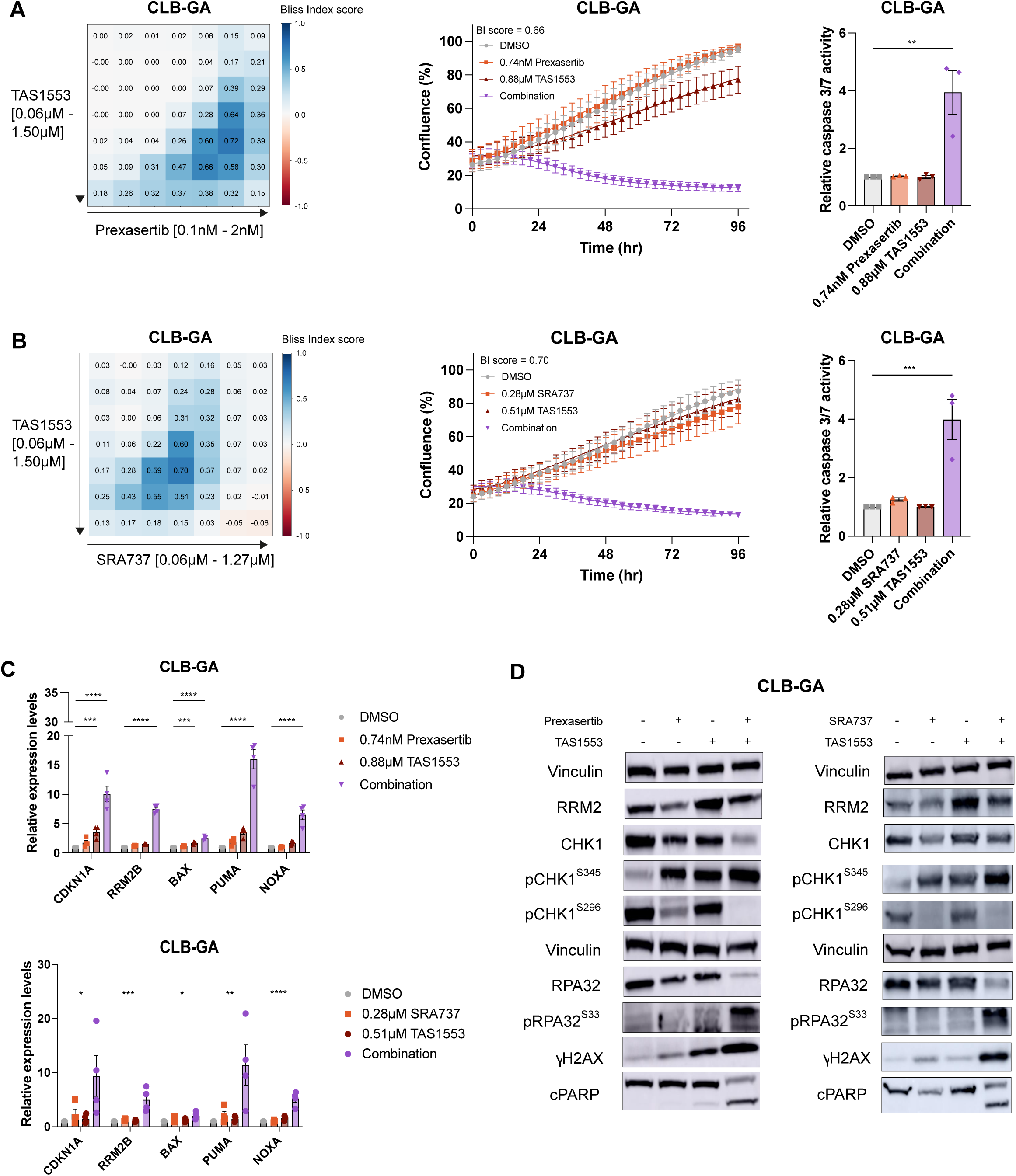
TAS1553 acts synergistically with prexasertib and SRA737 in CLB-GA. A. Heatmap showing the BI scores for CLB-GA after 96h of treatment with a range of concentrations of TAS1553 and prexasertib. Biological replicates= 4 in duplicate (*left*). Cell confluency was monitored for 96h and cells were treated with the concentrations with the highest BI score (0.74nM prexasertib and 0.88µM TAS1553). Biological replicates= 4 in duplicate, error bars represent SEM (*middle*). Caspase-Glo luminescence-based measurements of caspase 3/7 activity in CLB-GA following treatment with TAS1553 and prexasertib for 48h. Biological replicates= 3 in triplicate, error bars represent SEM. Significance was measured using the one-way ANOVA followed by a Dunnett’s multiple comparison test (**= p≤0.01) (*right*). B. Heatmap showing the BI scores for CLB-GA after 96h of treatment with a range of concentrations of TAS1553 and SRA737. Biological replicates= 4 in duplicate (*left*). Cell confluency was monitored for 96h and cells were treated with the concentrations with the highest BI score (0.28µM SRA737 and 0.51µM TAS1553). Biological replicates= 4 in duplicate, error bars represent SEM (*middle*). Caspase-Glo luminescence-based measurements of caspase 3/7 activity in CLB-GA following treatment with TAS1553 and SRA737 for 48h. Biological replicates= 3 in triplicate, error bars represent SEM. Significance was measured using the one-way ANOVA followed by a Dunnett’s multiple comparison test (***= p≤0.001) (*right*). C. RT-qPCR based measurements of mRNA expression of apoptotic genes *BAX, PUMA* and *NOXA*, and the *p21* target genes *CDKN1A* and *RRM2B* in CLB-GA after treatment with TAS1553 in combination with prexasertib (*top*) or SRA737 (*bottom*) that resulted in the highest synergy scores at 48h. Biological replicates= 4 in duplicate. Significance was measured using the one-way ANOVA followed by a Dunnett’s multiple comparison test (*= p≤0.05, ***= p≤0.001, ****= p≤0.0001), error bars represent SEM. D. Representative immunoblotting data for various replicative stress and DNA damage markers in CLB-GA upon treatment with TAS1553 in combination with prexasertib (*left*) or SRA737 (*right*) for 48h. The quantification of all replicates is shown in Supplementary Figure S5A.

**Figure S4:**
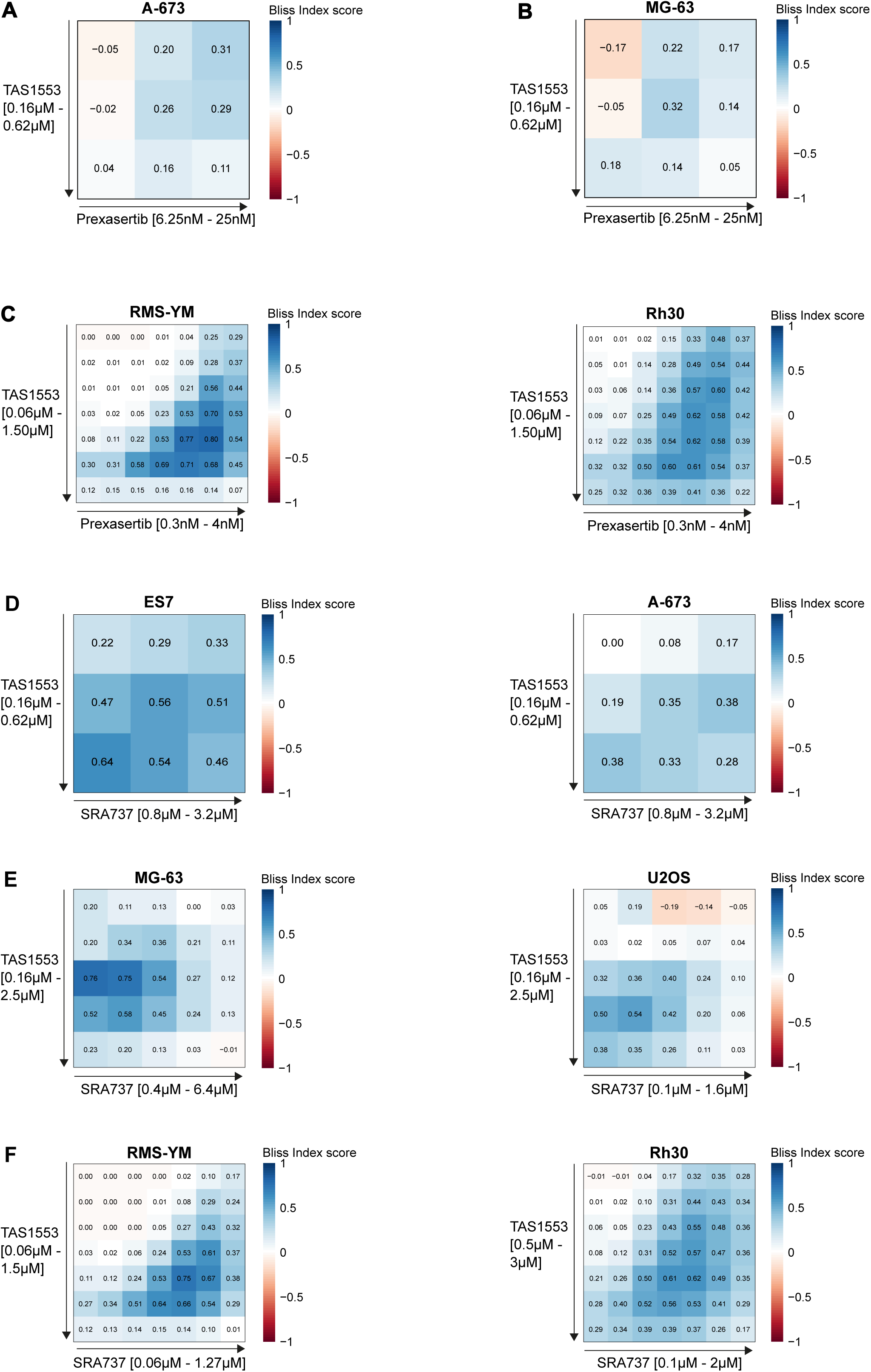
The combination of TAS1553 with prexasertib or SRA737 is synergistic in pediatric sarcoma cells *in vitro*. A. Heatmaps showing the BI scores for the Ewing sarcoma cell line A-673 after 72h of treatment with a range of concentrations of prexasertib and TAS1553. Biological replicates= 4 in triplicate. B. Heatmaps showing the BI scores for the osteosarcoma cell line MG-63 after 72h of treatment with a range of concentrations of prexasertib and TAS1553. Biological replicates= 4 in triplicate. C. Heatmaps showing the BI scores for the rhabdomyosarcoma cell lines RMS-YM (*left*) and Rh30 (*right*) after 72h of treatment with a range of concentrations of prexasertib and TAS1553. Biological replicates= 3 in duplicate. D. Heatmaps showing the BI scores for the Ewing sarcoma cell lines ES7 (*left*) and A-673 (*right*) after 72h of treatment with a range of concentrations of SRA737 and TAS1553. Biological replicates= 4 in triplicate. E. Heatmaps showing the BI scores for the osteosarcoma cell lines MG-63 (*left*) and U2OS (*right*) after 72h of treatment with a range of concentrations of SRA737 and TAS1553. Biological replicates= 4 in triplicate. F. Heatmaps showing the BI scores for the rhabdomyosarcoma cell lines RMS-YM (*left*) and Rh30 (*right*) after 72h of treatment with a range of concentrations of SRA737 and TAS1553. Biological replicates= 3 in duplicate.

**Figure S5:**
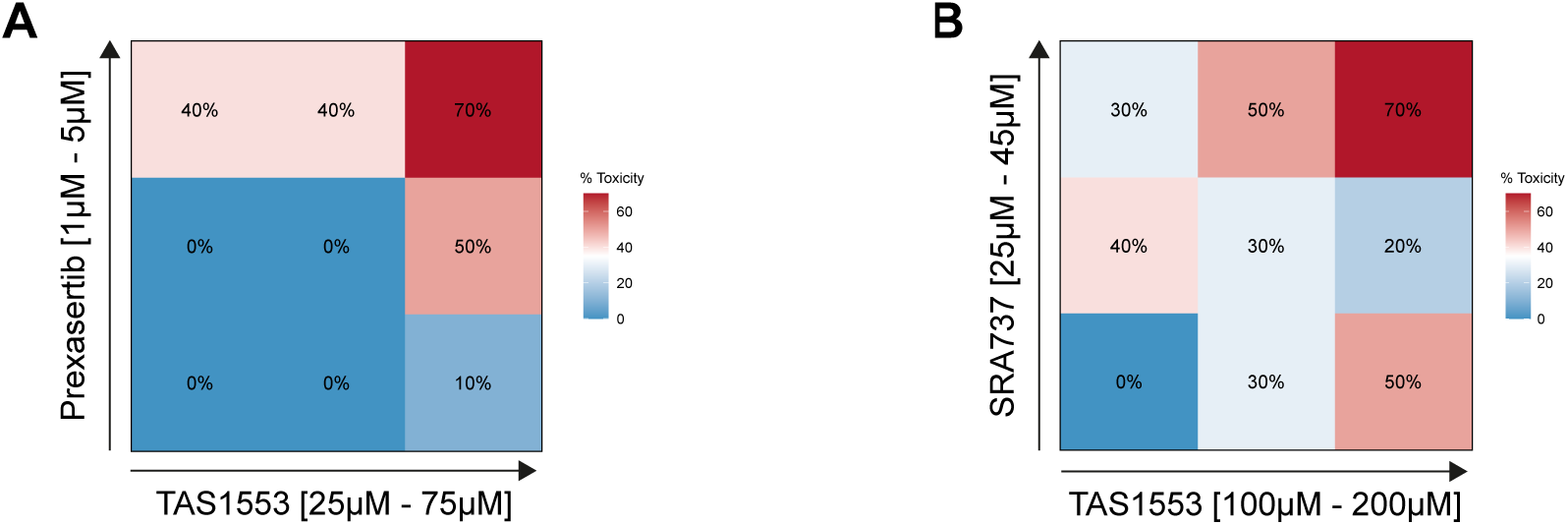
Determination of the MNTD for TAS1553 in combination with prexasertib or SRA737 in zebrafish. A. Determination of the MNTD for TAS1553 in combination with prexasertib in wildtype zebrafish embryos of 3 days-post-fertilization (dpf). The percentage of fish without toxicities are indicated in the boxes. B. Determination of the MNTD for TAS1553 in combination with SRA737 in wildtype zebrafish embryos of 3 dpf. The percentage of fish without toxicities are indicated in the boxes.

**Figure S6:**
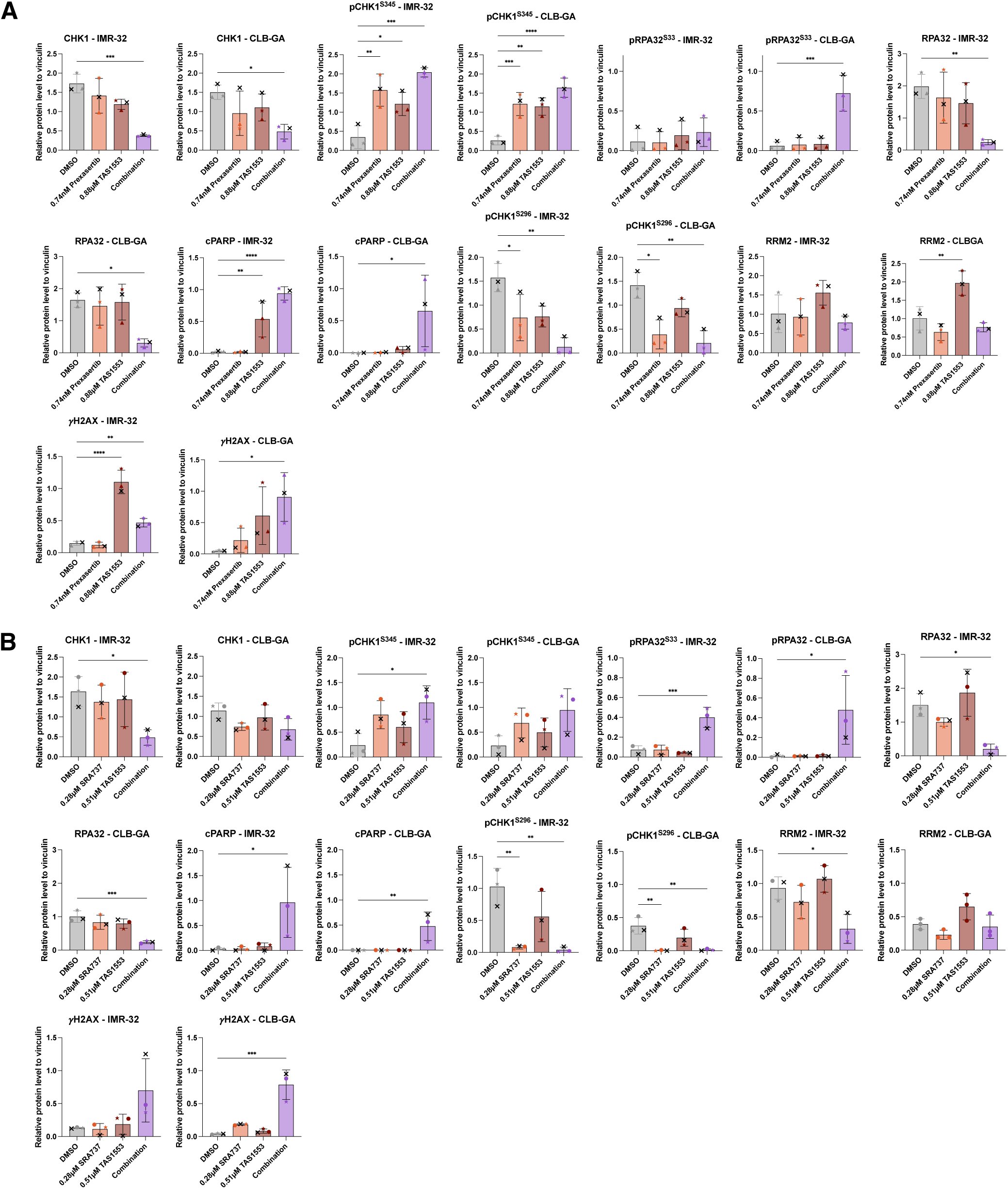
Quantification of the western blot data as displayed in. **Figure 3 and 4 and Appendix Figure S3.** A. Quantification of the immunoblotting of IMR-32 and CLB-GA treated with 0.74nM prexasertib and 0.88µM TAS1553 for 48h. Biological replicates= 3. Significance was measured using the one-way ANOVA followed by a Dunnett’s multiple comparison test (*= p≤0.05, **= p≤0.01, ***= p≤0.001, ****= p≤0.0001), error bars represent standard deviation (SD). B. Quantification of the immunoblotting of IMR-32 and CLB-GA treated with 0.28µM SRA737 and 0.51µM TAS1553 for 48h. Biological replicates= 3. Significance was measured using the one-way ANOVA followed by a Dunnett’s multiple comparison test (*= p≤0.05, **= p≤0.01, ***= p≤0.001), error bars represent SD.

**Figure S7:**
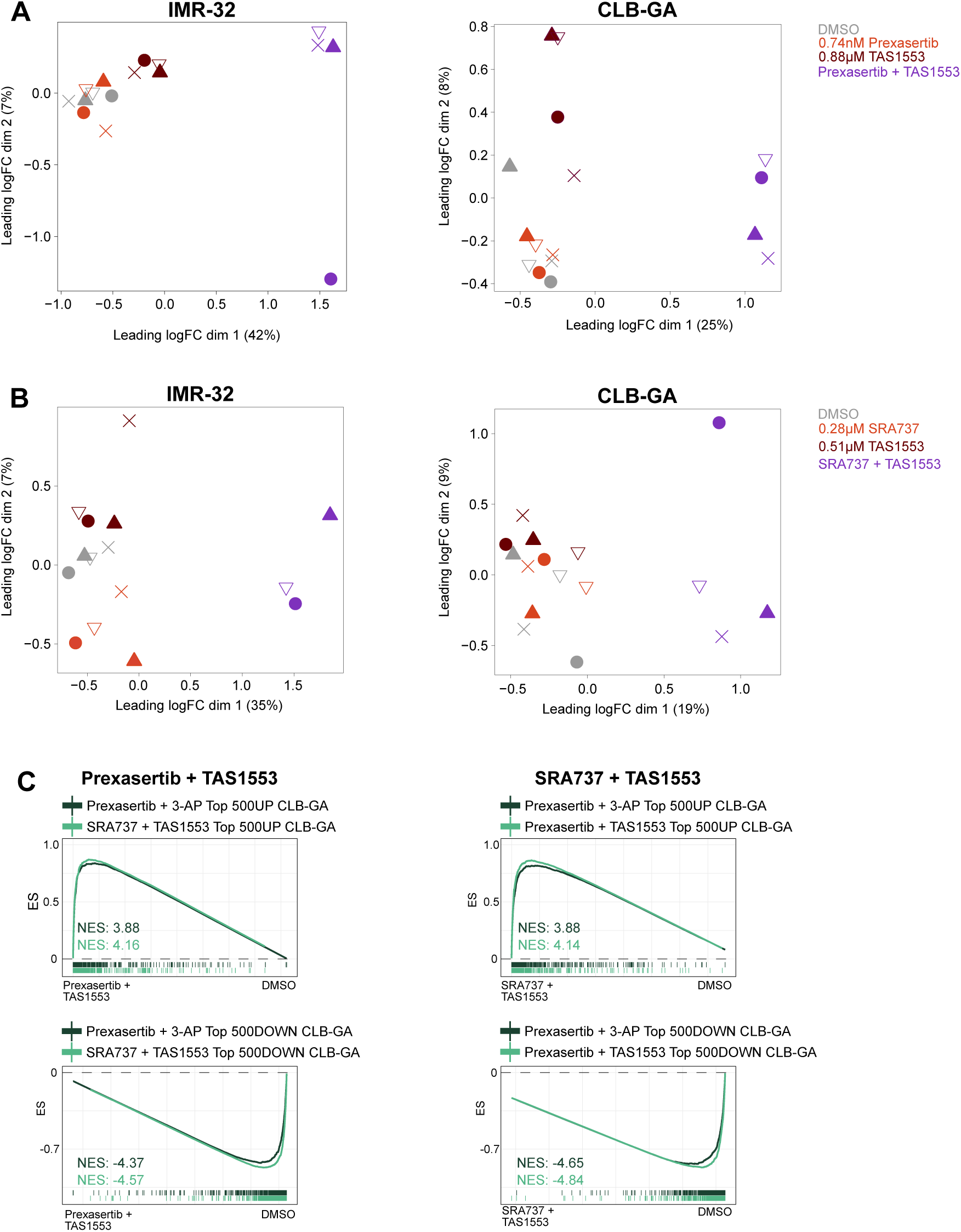
Gene expression signatures following TAS1553 exposure in combination with either SRA737 or prexasertib show significant overlap (*continued*). A. PCA showing the distinction between IMR-32 (*left*) and CLB-GA (*right*) cells treated with prexasertib (0.74nM) and TAS1553 (0.88µM) compared to cells treated with the single compounds or control (DMSO). Symbols represent different replicates. B. PCA showing the distinction between IMR-32 (*left*) and CLB-GA (*right*) cells treated with SRA737 (0.28µM) and TAS1553 (0.51µM) compared to cells treated with the single compounds or control (DMSO). Symbols represent different replicates. C. GSEA of the RNA-seq based transcriptome profiles following combined drug treatment with TAS1553 and prexasertib (*left*) or TAS1553 and SRA737 (*right*) in CLB-GA cells showing a significant overlap between genes enriched upon combination treatment of TAS1553 with prexasertib, TAS1553 with SRA737 or 3-AP with prexasertib. ES= enrichment score.

**Figure S8:**
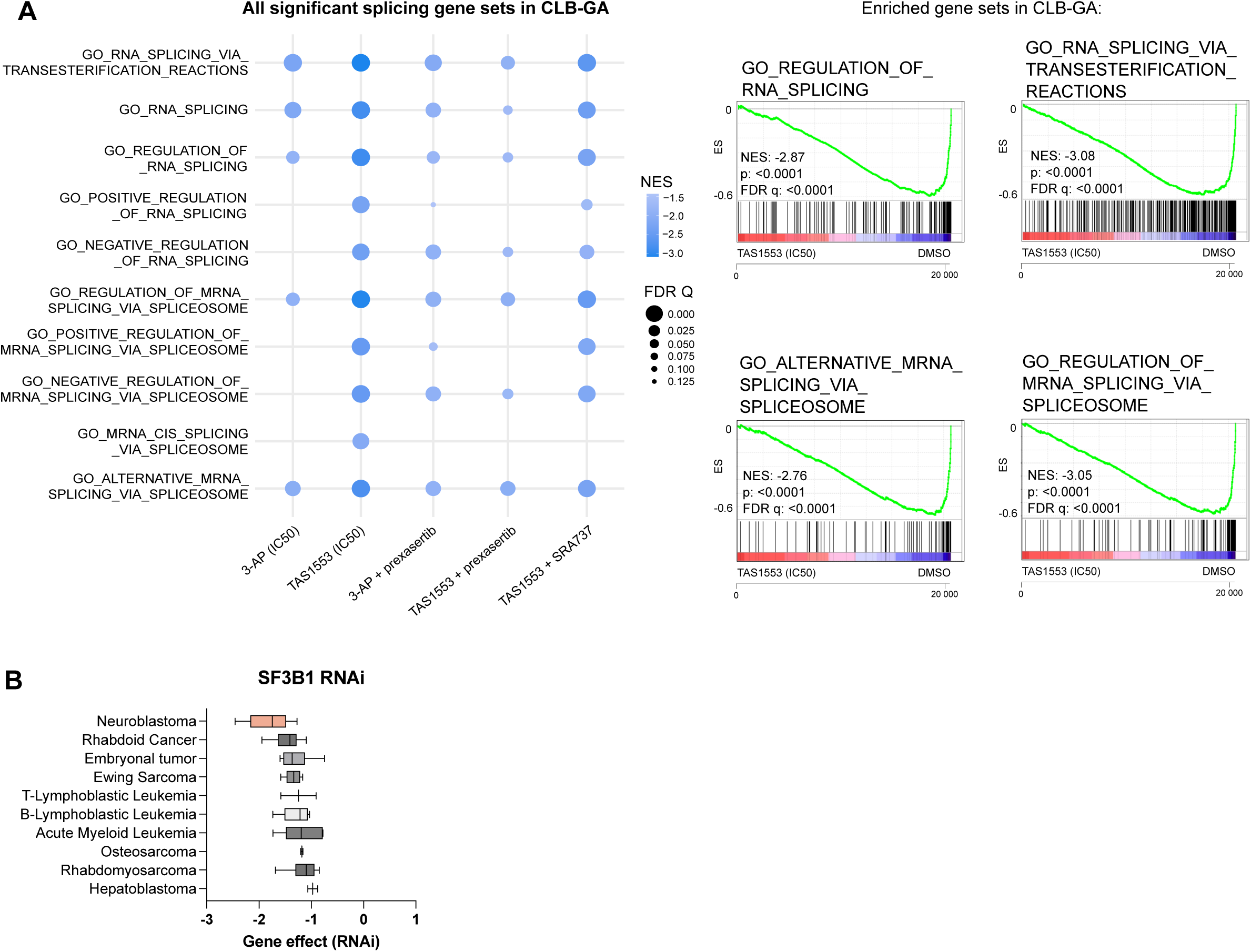
TAS1553 induces downregulation of gene sets involved in RNA splicing in neuroblastoma. A. Bubble plot showing the downregulated gene sets in CLB-GA involved in RNA splicing, downregulated after treatment with TAS1553 or one of the combinations with TAS1553 (*left*). Corresponding GSEA of the top two gene sets involved in RNA splicing or the spliceosome upon treatment with the IC_50_ of TAS1553 (*right*). B. DepMap analysis (https://depmap.org/portal) of RNAi screen of SF3B1 in pediatric cancers indicating high dependency in neuroblastoma.

